# Maternal opioid use with and without hepatitis C infection disrupts the structure and immune landscape of the maternal-fetal interface

**DOI:** 10.1101/2025.07.30.667651

**Authors:** Heather E. True, Brianna M. Doratt, Sheridan Wagner, Delphine C. Malherbe, Nathan R. Shelman, Mahdi Eskandarian Boroujeni, Cynthia Cockerham, John O’Brien, Ilhem Messaoudi

## Abstract

Maternal opioid use disorder (OUD) poses significant risks to maternal and fetal health. Adverse outcomes associated with maternal OUD are believed to be mediated, in part, by changes in placenta structure and function; however, few studies have addressed this question. Here, we utilized a combination of flow cytometry, histology, spatial and single-cell transcriptomics to uncover the impact of OUD on placental tissues. Given that nearly half of subjects with chronic OUD contract hepatitis C (HCV), we further stratified our findings by maternal HCV status. Our results indicate that maternal OUD leads to a higher incidence of vascular malperfusion accompanied by increased levels of inflammatory markers and dysregulated secretion of placental development factors. Furthermore, spatial transcriptomics revealed that maternal OUD disrupts the communication between trophoblasts and immune cells important for placental vascular development. Additionally, CellChat analysis revealed aberrant VEGF and FN1 signaling across trophoblast, endothelial, and myeloid cells. Processes associated with tissue homeostasis and repair were also downregulated across trophoblast and leukocytes. Frequencies and responses to ex-vivo stimulation of decidual macrophages and cytolytic NK cells, critical for tissue remodeling and fetal tolerance, were decreased. Finally, transcriptional analyses of placental leukocytes also indicate shifts towards more regulatory/tissue surveillant phenotypes. Altogether, these results highlight the significant disruptions to placental health by maternal OUD.

**One Sentence Summary:** Maternal opioid use disorder ± hepatitis C coinfection disrupts placental structure, immune function, and cell-cell communication.

## INTRODUCTION

Opioid use disorder (OUD) continues to be a significant public health concern, with national overdose deaths nearly quadrupling from 2012-2022 (*1*). Drug overdose mortality rates in pregnancy have nearly doubled from 2018 to 2021(*2*), and OUD in pregnancy has been linked to adverse effects for both pregnant women and fetuses. For the pregnant woman, OUD has been associated with higher rates of mortality, placental abruption, preterm labor, and oligohydramnios (*3, 4*). For the offspring, reported complications include increased risk of death, poor fetal growth, preterm birth, birth defects, and impaired neurodevelopment (*5–8*).

These adverse outcomes are believed to be mediated by alterations in placental vascular development and function (*9, 10*). The placenta is a transient fetal-maternal organ that provides a conduit for nutrient, gas, and waste exchange between maternal and fetal circulation (*11*). It is comprised of maternal- and fetal-derived components, termed the decidua basalis and chorionic villous, respectively (*12*). The decidua basalis gradually thins as the fetal chorionic villous takes over supporting the developing fetus (13). However, the decidua maintains an essential role throughout pregnancy by providing a scaffolding for the growing placenta and fetus, producing prostaglandins and other inflammatory factors to initiate contractions (*14*). The chorionic villous is the main site for angiogenesis, where the development of new capillaries create a network with maternal circulation (*15*). Dense networks of villi increase the surface area available for exchange of gasses and nutrients between the chorion and the maternal uterine lining (*16*). Therefore, the placenta is a key mediator of fetal/maternal interaction by providing signals that regulate fetal growth and tolerance (*17*).

A healthy placenta is composed of trophoblast, endothelial, and stromal cells that facilitate fetal organ development, placental circulation, and hormone/cytokine signaling (14). Trophoblast cells are specialized fetal epithelial cells that differentiate from the outer layer of the developing embryo and maintain placental integrity as well as mediate the complex interplay with the maternal immune system (*12*). Placental endothelial cells maintain the integrity of the placental barrier by signaling to trophoblasts, stromal cells, and immune cells to promote vascular remodeling, optimize maternal blood flow, and antiviral defenses (*18, 19*). Finally, placental stromal cells have potent immunosuppressive properties that prevent immune-mediated rejection of the fetus (*17, 20*). In addition, the decidua is composed of maternally derived immune cells including T cells, NK cells, macrophages, and few dendritic cells (*21*). Decidual immune cells play critical roles in maintaining immune tolerance (*22*), promoting fetal growth (*23*), and responding to infections at the maternal-fetal interface (*24, 25*).

Immune cells of the chorionic villous are comprised exclusively of primitive, fetal yolk sac-derived macrophages called Hofbauer cells (HBC) and infiltrating placenta-associated maternal monocytes and macrophages (PAMM). HBCs are critical for placental morphogenesis, maintaining fetal tolerance, and protection of the fetus from maternal infection and inflammation (*26, 27*). PAMMs, However, are a heterogenous population of macrophages. PAMM1A cells are the most abundant of the PAMMs and are morphologically similar to macrophages and adhere to sites of placental injury, secrete tissue repair factors, and regulate trophoblast function (*28*). PAMM1B cells are monocytes with unique gene signatures from circulating blood monocytes and are thought to be PAMM1A precursors (*28*). Finally, PAMM2 macrophages are present in low abundance in healthy pregnancies and are considered contaminating macrophages from the maternal decidua (*28*).

Few studies have addressed the impact of opioids on placental structure and immune function (*29, 30*). Methadone, an opioid agonist, and buprenorphine, a partial opioid agonist, are commonly used therapeutics in the treatment of OUD. Pregnant individuals treated with opioid maintenance therapy are at a significantly higher risk for placental dysmaturity, characterized by irregular villi, decreased syncytial knots, hypervascularity, and villous edema (*30, 31*). Additionally, pregnant individuals who receive prescription opioids during pregnancy (regardless of gestational age) are at a higher risk of placental insufficiency and/or ischemia, compared to healthy pregnancies (*32*). However, the mechanisms by which opioids result in these adverse outcomes remain poorly understood. Furthermore, pregnancies exposed to opioids are often further challenged by maternal hepatitis C (HCV) infection, as over half of people who inject drugs develop HCV within five years (*33*). Additional studies are necessary to uncover the impact of maternal OUD ± HCV infection on placental structure and immunity.

Thus, we collected maternal (decidua basalis) and fetal (chorionic villous) placental tissues from healthy pregnant participants and those with severe OUD with and without HCV infection (± HCV) to infer the impact of OUD±HCV on the structure, function, and immune landscape of the maternal-fetal interface. Briefly, maternal plasma and supernatants from tissue homogenate were assayed for immune mediators. Isolated leukocytes from each compartment were subjected to a series of assays to uncover the phenotypic, functional, and transcriptional landscape of placental immune cells.

Additional H&E-stained placental tissues were reviewed for histological pathologies and further interrogated by Visium spatial transcriptomics. We next inferred intercellular communication networks from our spatial transcriptomics dataset using CellChat (*34*).

Finally, given the high prevalence of HCV infection in this cohort, we stratified our findings by maternal HCV status.

## RESULTS

### Subhead 1: Maternal Opioid Use and HCV infection lead to adverse maternal-fetal outcomes

Decidua and chorionic villous tissues were isolated from placentas of full-term pregnancies (>37 weeks) of participants ± OUD (Fig. 1A). Maternal demographics are summarized in Table 1. At enrollment, all participants in the OUD group were screened per the Diagnostic and Statistical Manual of Mental Disorders, Fifth Edition (DSM-V) criteria for severe OUD, as defined by meeting 4+ criteria at enrollment (Table 1). All participants in the OUD group reported a history of illicit opioid use (heroin, fentanyl, and/or non-prescribed opiates, buprenorphine, and methadone products) with 39.5% of participants using more than one substance (Fig. 1B). We then stratified participants based on their HCV status, using self-reporting and confirmed by HCV IgG antibody detection and RTq-PCR at delivery (Table 1). Of the total 96 participants with OUD screened for HCV, 46 participants were HCV-IgG positive indicating history of infection and 18 had a viral load detected by RTq-PCR indicating active HCV infection at delivery (Table 1). On average, participants in the OUD_HCV+ group had slightly longer gestations (Table 1). Other maternal factors including maternal age, pre-pregnancy body mass index (BMI), delivery mode, and fetal sex were comparable (Table 1).

**Fig. 1:**
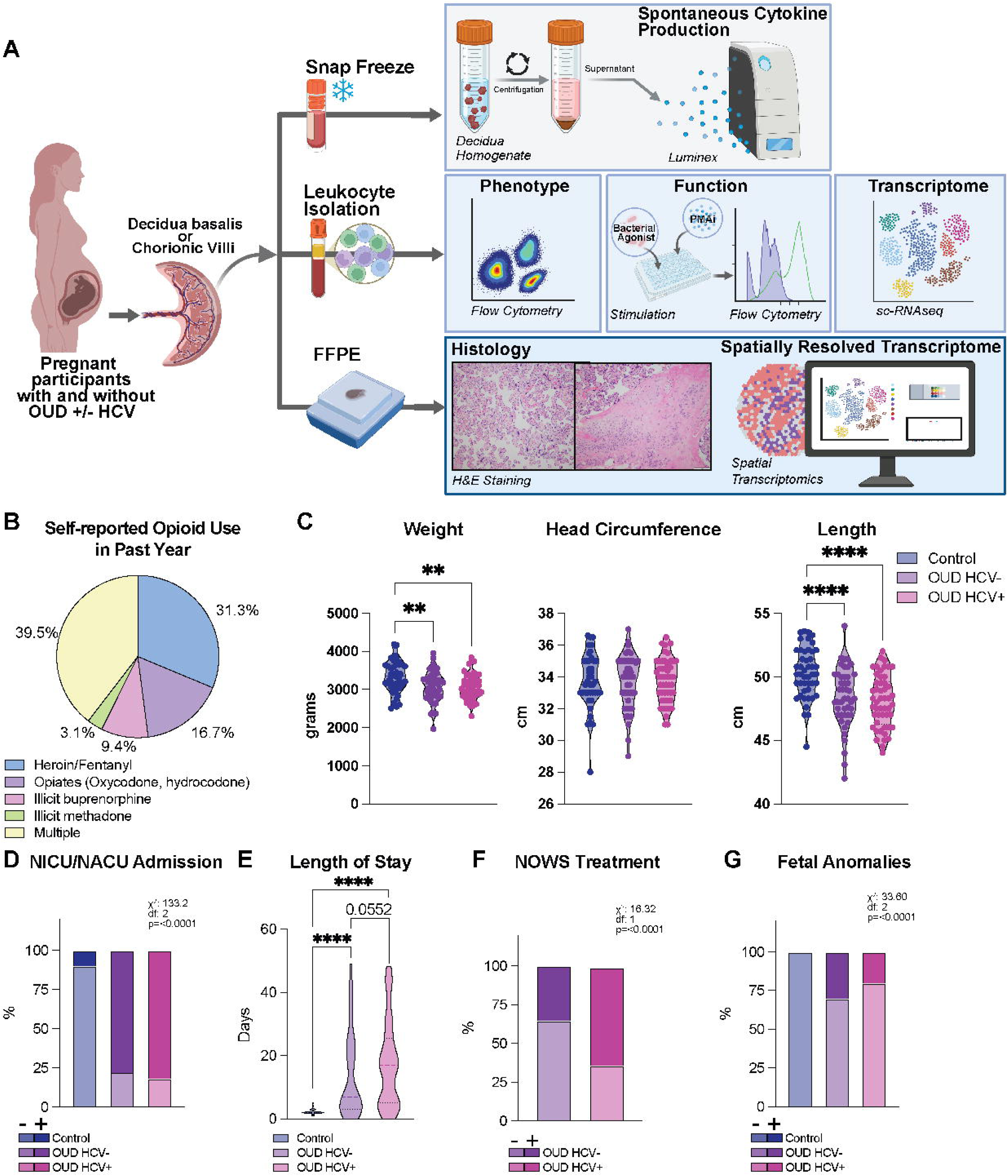
Maternal OUD±HCV leads to adverse maternal-fetal outcomes. A) Experimental design. B) Pie chart depicting self-reported substance use at enrollment over the previous 1 year. “Multiple” indicates the use of 2 or more of the listed substances. C) Violin plots of newborn birth weight (g), head circumference (cm), and length (cm). D) Stacked bar plot depicting the % of newborns admitted to intensive care units (NICU/NACU). E) Violin plot of newborn length of stay in intensive care. (F-G) Stacked bar plots showing the % of F) newborns that received pharmacological interventions for the treatment of Neonatal Opioid Withdrawal Syndrome (NOWS), and G) newborns presenting with fetal anomalies. **=p<0.01, ****=p<0.0001.

**Table 1:**
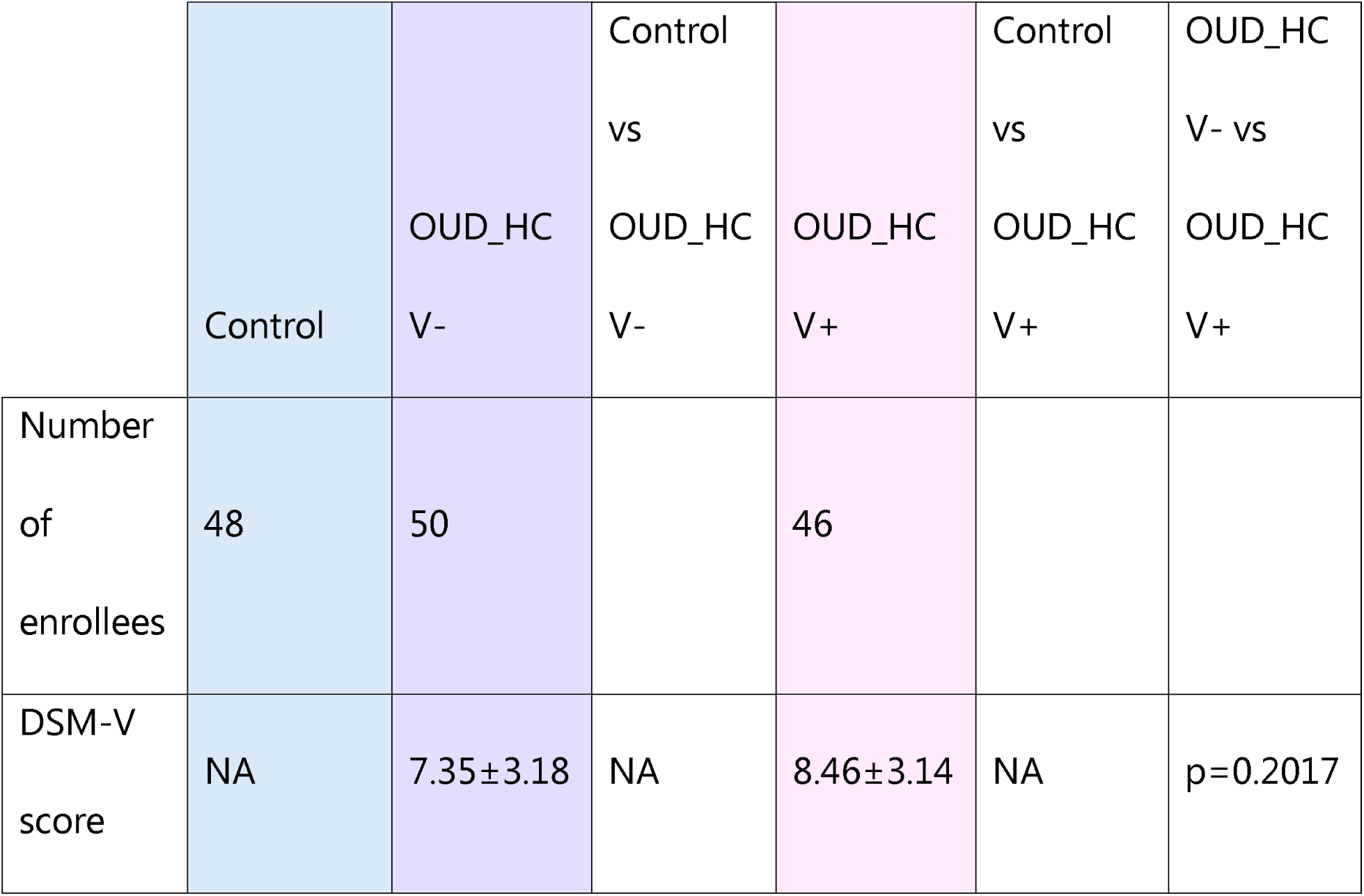
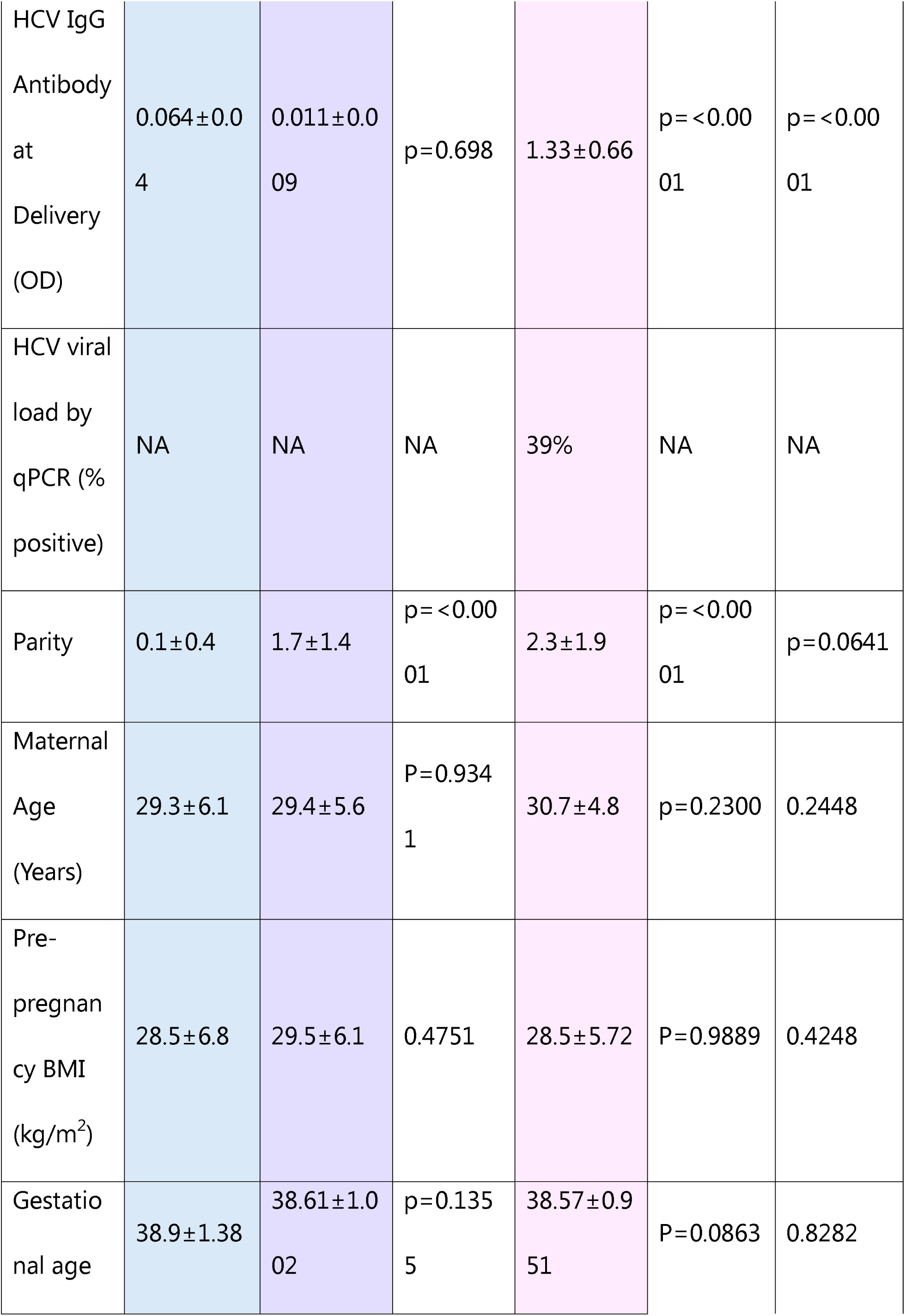
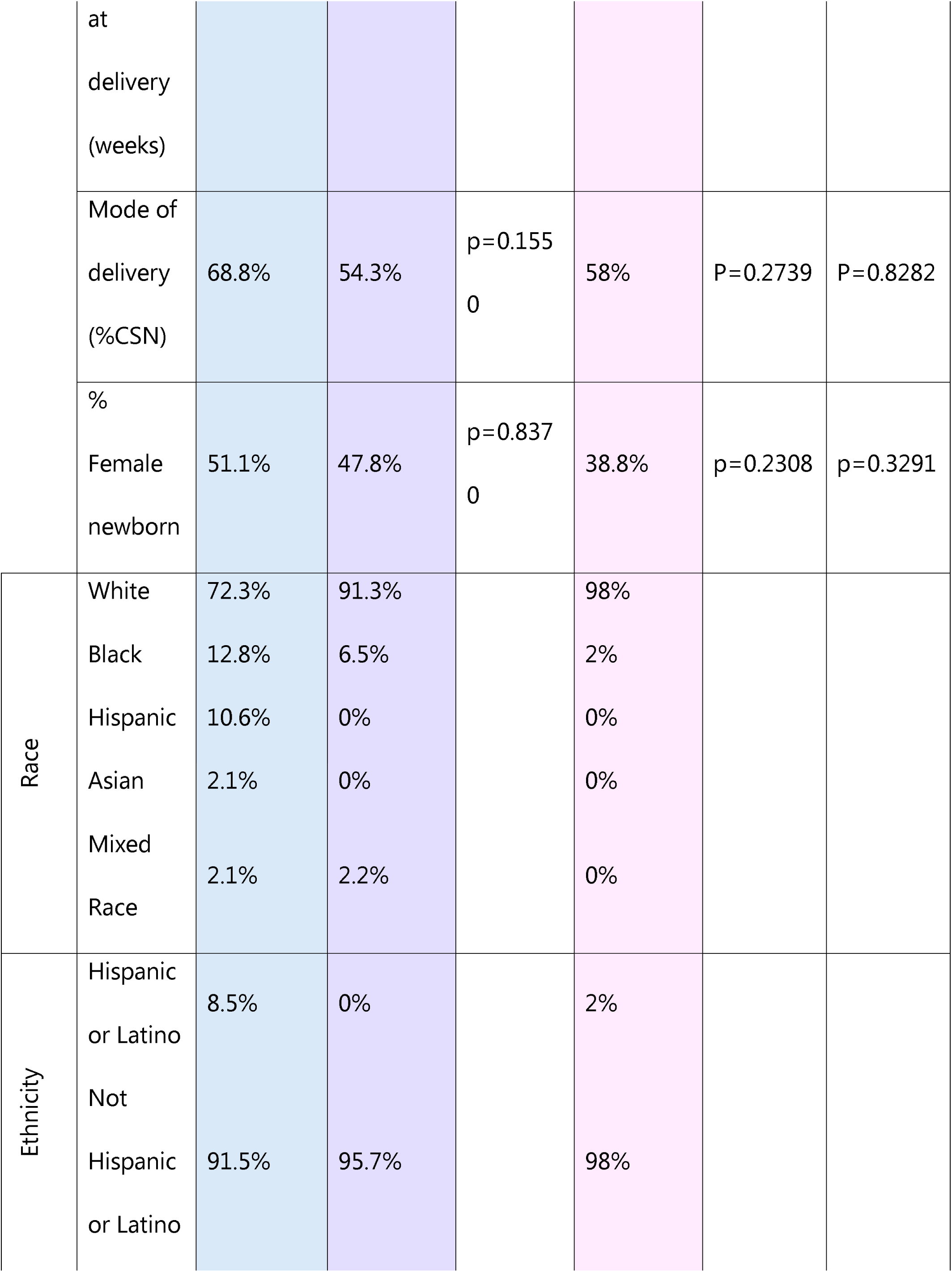
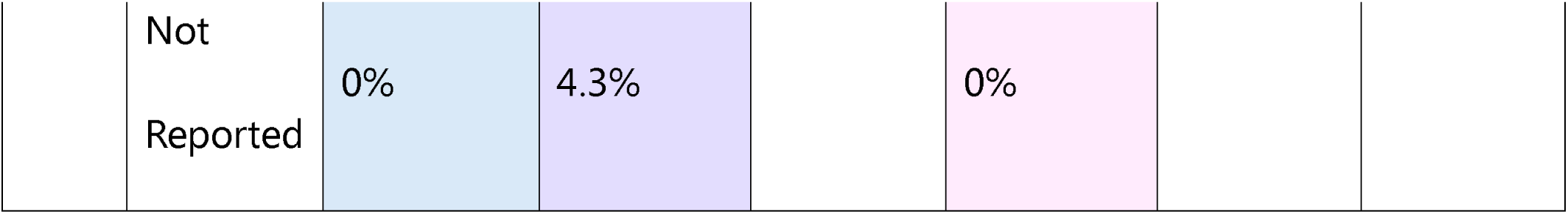
Cohort Characteristics.

Maternal OUD exerts acute and long-term adverse outcomes for newborns, including growth restriction, developmental delays, and increased likelihood of chronic diseases (*4*). In line with these observations, OUD was associated with smaller newborns (Fig. 1C) who were more likely to be admitted to neonatal intensive care units (NICU) (Fig. 1D) for a protracted length of time (Fig. 1E). These findings were exacerbated by maternal HCV infection (Fig. 1D-E), as was the need for pharmacological treatment (morphine) for neonatal opioid withdrawal syndrome (NOWS) (Fig. 1F). Finally, fetal anomalies (birth defects) were more abundant in newborns of participants with OUD without HCV infection (Fig. 1G). Overall, these findings highlight the profound impact of maternal OUD on fetal outcomes that is further exacerbated by HCV infection.

### Subhead 2: Altered inflammatory milieu and immune profiles of the decidua basalis with maternal OUD±HCV

The placenta undergoes dynamic changes throughout gestation to support fetal growth while simultaneously preventing fetal rejection (*35, 36*). These changes are highly regulated by immune mediators and growth factors (*36*). Therefore, we measured immune mediators and angiogenesis factors in decidua tissue homogenate. We detected a significant increase in pro-inflammatory cytokines (TNFα, IL-6RA, IL-1Ra, IL-18, IL-2), chemokines (IP-10, MIP-1β, CXCL9, IL-8), and EGF with OUD (Fig. 2A). However, levels of TH2 cytokine IL-5 and angiogenesis factors (FLT-1/4) were reduced compared to control (Fig. 2A). We then carried out a sparse partial least squares differential analysis (sPLSDA) to identify factors that separate the three groups (Fig. 2B-C). Factors including IL-4, PECAM1, and IL-22 delineated the OUD_HCV-group while those that differentiate the OUD_HCV+ group include TNFβ and VEGFR2 (Fig. 2C). Overall, these findings indicate that maternal OUD is associated with increased inflammation in the decidua that is further amplified by HCV coinfection.

**Fig. 2:**
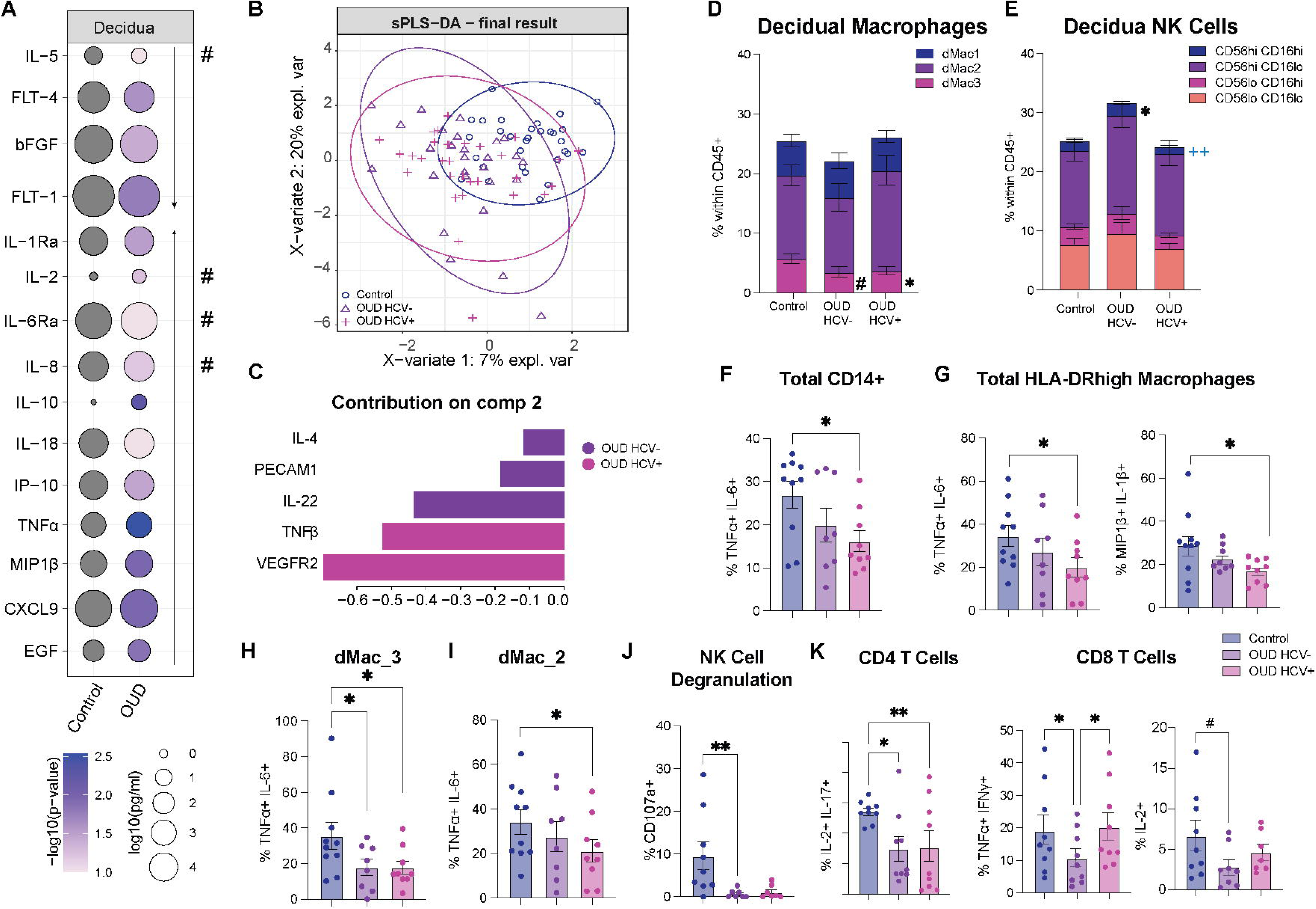
Decidual immune cell phenotype and function are altered by maternal OUD±HCV. A) Bubble plot of factors detected in the supernatant of decidua tissue homogenate. The size of the bubble represents the concentration (pg/ml) where the color represents the significance (p-value). B) Principle Component Analysis (PCA) from the sPLSDA comparisons of control, OUD_HCV-, and OUD_HCV+ groups by concentration of markers measured in decidua tissue homogenate displayed in 1-2 space. C) Bar plot of markers delineated by group from the sPLS-DA component 1. (D-E) Stacked bar plots of D) % HLA-DRhigh decidual macrophage subsets, and E) % NK cell subsets within the CD45+ population. (F-H) Bar plots of % response of F) total CD14+ cells, G) total HLA-DRhigh macrophages, H) dMac_3, and I) dMac_2 cells to bacterial TLR ligand stimulation. J) Bar plot of the % of NK cells expressing CD107a in response to PMAi stimulation. K) Bar plot of % response of CD4+ (left) and CD8+ (right) T cells to PMAi stimulation. Symbols in black denote comparisons to control whereas blue symbols denote significance between OUD_HCV- and OUD_HCV+ groups. #=p<0.01, *=p<0.05, **^/++^=p<0.01.

Furthermore, it has been well established that markers of placental dysfunction can be detected in maternal circulation (*37*). Therefore, we measured levels of placental development in maternal plasma. We first identified significantly different markers between control and total OUD groups and revealed lower levels of PDGF-AA, FGF1, Angiopoetin-1, ANGPTL-4/6, and FLT-1 with maternal OUD (Fig. S1A). Decreased levels of these markers in maternal plasma generally reflect impaired placental vascular development and reduced fetal nutrient and oxygen supply (*38*). Furthermore, we detected elevated levels of PAPP-A, PECAM-1, FLT-4, MMP-8, and GM-CSF in maternal plasma in the OUD group versus control (Fig. S1A). High levels of these markers have been associated with placental dysfunction and hypertensive disorders in pregnancy (*39*). Further sPLSDA analysis delineating the OUD_HCV+ group include Tenascin C, EGF, FGF-1, IL-6RA, and PDGF-AA, while those that differentiated the OUD_HCV-group include c-kit, angiopoietin-1, FLK, and MMP8 (Fig. S1C). However, PECAM1, MMP9, HGF, FLT4, ANGPTL3/4/6, and FLT1 levels segregated the control from the OUD groups (Fig. S1C). Altogether, these findings highlight that maternal OUD leads to a pro-inflammatory milieu and elevated placental repair signaling.

We next assessed the decidual immune cell compartment using flow cytometry (Fig. S1D). While no significant differences in the abundance of total monocytes, dendritic, NK, or T cells were observed (Fig. S1E), we report significant shifts in the frequency of HLA-DR^high^ decidual macrophage and NK cell subsets with maternal OUD±HCV (Fig. 2D-E). Specifically, the frequency of dMac_3 macrophages (FOLR2^high^S100A8/9^low^CD9^-^), that are mature decidual macrophages important for tissue remodeling and clearance of cell debris in late pregnancy (*21*), was reduced with maternal OUD (Fig. 2D). Furthermore, we report a decrease in cytolytic CD56^high^CD16^high^ NK cells, recently described as critical for regulating immune responses and supporting placentation in healthy pregnancies (*40*), in OUD_HCV+ subjects (Fig. 2E).

To evaluate differences in the functional capacity, we next assayed responses to *ex vivo* stimulation (Fig. 1A). Decreased responses to bacterial ligands by total decidual macrophages was noted with maternal OUD and HCV infection (Fig. 2F), driven by dampened responses generated by HLA-DRhigh macrophages (Fig. 2G), namely dMac_3 and dMac_2 subsets (Fig. 2H-I). Additionally, degranulation by dNK in response to stimulation by PMA/ionomycin (PMAi) was decreased with maternal OUD (Fig. 2J). Finally, although no differences were observed in the abundance of decidual CD4 or CD8 T cells (Fig. S1E), T cell responses to polyclonal stimulation by PMAi were also decreased with maternal OUD (Fig. 2K). Altogether, these findings indicate that maternal OUD±HCV alters the abundance and function of immune cells in the decidua.

### Subhead 3: Altered inflammatory milieu and immune profiles of chorionic villous (fetal) placental tissues with maternal OUD±HCV

Next, we compared differences in inflammatory and angiogenesis factors in villous tissue homogenate between control and total OUD groups (Fig. S2A). Levels of IL-12p70 and IL-7 were increased while those of IL-6 and IL-8 were decreased with maternal OUD (Fig. S2A). Changes in levels of these cytokines have been associated with pre-eclampsia and preterm birth (*41, 42*). We next carried out a sPLSDA analysis to tease out differences between all three groups (Fig. S2B-C). Notable markers that delineated the OUD_HCV+ group include TNFβ, IFNγ, GM-CSF, IFNα-2, and Eotaxin, whereas IL-17A, IL-12p70, IL-7, IL-13, and IL-4 delineated the OUD_HCV-group (Fig. S1B). However, IL-10, TNFα, EGF, and IL-1α levels segregated the control group from the OUD groups (Fig. S1B). Altogether, these findings highlight that maternal OUD±HCV shifts the placental milieu towards a hyperinflammatory state.

We then used flow cytometry to characterize the immune landscape of the chorionic villous (Fig. 1A, Fig. S2D). No differences were found in the abundance of HBCs with maternal OUD regardless of HCV status (Fig. S2E). The frequency of PAMM1A cells increased in the OUD_HCV+ group compared to controls and OUD_HCV-, while that of PAMM2 cells decreased (Fig. S2E). A modest decrease in the abundance of PAMM1B cells in the OUD_HCV+ group, relative to OUD_HCV-was seen (Fig. S2E). No differences were observed in the phagocytosis potential (Fig. S2F) or cytokine response of chorionic villous macrophages following stimulation with bacterial TLR ligands (Fig. S2G). These findings indicate that maternal OUD and HCV infection has a more profound impact on the inflammatory milieu and immune cell landscape of the decidua relative to villous tissues.

### Subhead 4: Maternal OUD rewires the transcriptional profile of placental immune cells

We next profiled the transcriptome of CD45^+^ FACS-sorted placental leukocytes from decidua and chorionic villous tissues using single-cell RNA sequencing (scRNA-seq) and integrated these datasets to construct a comprehensive transcriptional profile of the total placental immune landscape (Fig. 1A). Our analysis revealed 17 unique immune cell clusters encompassing previously defined populations from term decidua (*21*) and chorionic villous (*43*) (Fig. 3A-B, Table S1). Specifically, we identified T cells (*CD3*^+^*IL7R*^+^*CD8A*^+^), four populations of decidual NK cells (dNK1:*NKG7*^+^, dNK2:*NKG7*^+^*GZMB*^high^*XCL1*^+^*CD160*^+^, dNK3:*NKG7*^+^*GZMB*^low^*CD160*^+^, and dNK_proliferating:*NKG7*^+^*MK67*^+^), and three B cell populations (B cell_naive: *MS4A1*^+^, B cell_memory: *MS4A1*^+^ *CD27*^+^, and B cell_inflammatory: *MS4A1*^+^*CD69*^+^*HLA-DRA*^+^) (Fig. 3A-B). Among innate immune cells, we delineated two dendritic cell populations (pDC:*CD38*^+^*GZMB*^+^, mDC:*CD1C*^+^*CD38*^+^*CD68*^+^); three *CD14*^+^*HLA-DR*^high^ decidual macrophage populations (dMac_1:*FCN1*^+^, dMac_2:*CD9*^+^*TREM2*^+^, dMac_3:*FOLR2*^+^*TREM2*^+^), as well as PAMM (PAMM1B: *HLA-DRA*^+^*S100A9*^+^*FCN1*^+^*SELL*^+^, PAMM1A:*HLA-DRA*^+^*CD9*^+^*CXCL9*^+^, PAMM2:*HLA-DRA*^+^*FOLR2*^+^) and HBC (*HLA-DRA*^low^*FOLR2*^+^*NFKBIZ*^-^) subsets (Fig. 3A-B). Our results indicate a marginal decrease in mDC and PAMM2 cells in the OUD_HCV-group compared to control, whereas frequencies of PAMM1B cells were expanded (Fig. 3C). These shifts were modulated by maternal HCV infection (Fig. 3C).

**Fig. 3:**
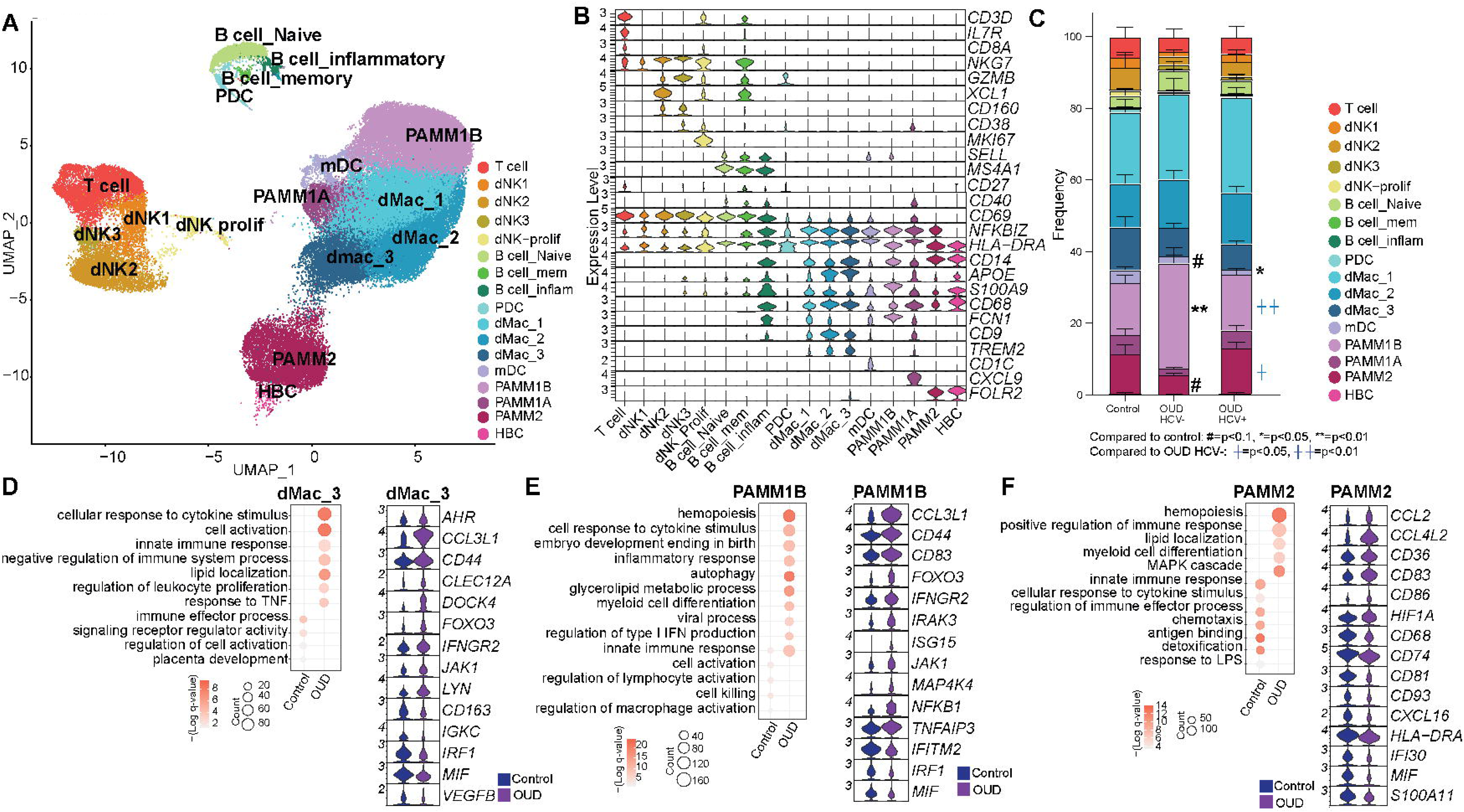
Maternal OUD impacts the transcriptional profile of placental immune cells. A) UMAP of cell subsets within placental tissue (73,256 cells). B) Violin plots showing expression levels of marker genes used for cluster identification. C) Stacked bar graph of cluster frequencies between groups. (Control: 15,069 cells, OUD_HCV-: 26,381 cells, OUD_HCV+: 31,806 cells). (D-F) Bubble plots of differentially expressed genes (DEGs) between control and OUD groups (left) and violin plots showing representative DEGs from the select GO terms (right) for the shown cluster. For the bubble plots, the size of the bubble represents the number of genes associated with GO term and the color represents the significance (q-value). ^#^=p<0.1, *^/†^=p<0.05, **^/††^=p<0.01).

We next identified differentially expressed genes (DEG) and performed functional enrichment to uncover the impact of maternal OUD and HCV infection on the placental immune landscape. First, we identified DEGs between control and the total OUD group, regardless of maternal HCV status. Most DEGs were detected in maternal-derived monocytes/macrophages (dMac and PAMM subsets). Genes upregulated in dMac_1 with OUD mapped to inflammatory responses and effector immune functions (*CCL4L2, CD109, CD86*) (Fig. S3A). However, genes downregulated with OUD were associated with regulation of immune activation (*FOS, JUN, MIF*) (Fig. S3A), suggesting aberrant activation of inflammatory dMac_1 cells with maternal OUD. Within dMac_2, genes associated with wound healing, antimicrobial responses, and hemopoiesis were upregulated with maternal OUD (*CD44, LYN, IFNAR2*) (Fig. S3B) indicating increased activation of dMac_2 cells with maternal OUD. Similarly, genes associated with responses to cytokines, leukocyte proliferation, and immune activation (*JAK1, CCL3L1, DOCK4, AHR*) were upregulated with OUD in dMac_3 (Fig. 3D). In contrast, genes associated with regulation of signaling receptor activity and placenta development were downregulated ( *IRF1, MIF, VEGF*) (Fig. 3D). Overall, these findings highlight that maternal OUD rewires decidual macrophages towards a hyperinflammatory and activated state.

In the PAMM1B subset, our analysis revealed increased gene expression associated with cytokine responses, embryo development, and autophagy (*IRAK3, IFNGR2, TNFAIP3*) while genes associated with antiviral responses (*IRF1, MIF, IFITM2*) were downregulated with maternal OUD (Fig. 3E). In the PAMM1A subset, genes associated with response to wounding, embryo development, and myeloid cell differentiation (*AHR, CCL20, CD9*) were upregulated while genes associated with antigen presentation and myeloid cell activation (*C1QA, CASP4, CD163*) were downregulated with maternal OUD (Fig. S3C). Finally, the PAMM2 subset showed an increased expression of genes involved in hemopoiesis, lipid localization, and myeloid cell differentiation (*CD36, CCL2, CD83*) in the OUD group (Fig. 3F) while downregulated genes mapped to immune effector processes, cell migration, and antigen binding (*CXCL16, HLA-DRA, IFI30*) with maternal OUD (Fig. 3F). Overall, these findings highlight that maternal OUD skews PAMM cells towards a more regulatory/tissue surveillant phenotype.

In the T cell cluster, downregulated genes associated with cytokine signaling, cell killing, lymphocyte activation, and inflammatory responses (*FABP5, HLA-DRB1, IFNG, LYST*) in te OUD group (Fig. S3D). Given that decidual T cells in late healthy pregnancy normally display a highly differentiated effector memory phenotype (*44*), these results suggest that maternal OUD may hinder labor progression. Decidual NK cells also become more activated and inflammatory in late pregnancy to facilitate labor onset (*45*). Genes upregulated in the cytotoxic dNK2 subset include genes involved in viral responses (*GZMB, IFI6, CCL3L1*) while downregulated genes were associated with apoptotic signaling pathways and unfolded protein response (*HSPA8, CACYBP, DUSP5*) in the OUD group (Fig. S3E). These findings indicate that maternal OUD leads to heightened activation and inflammation of cytotoxic NK cells that are poised towards antiviral responses. Lastly, in the mDC cluster, upregulated genes mapped to hemopoiesis and cytokine responses (*CCL4L2, CD86, CXCR4*), while those associated with antigen presentation were decreased (*GNAS, HLA-DQA2, IRF1*) with maternal OUD (Fig. S3F). These results suggest that mDC function is skewed towards a heightened activation state but defective antimicrobial responses.

### Subhead 5: Maternal HCV infection shifts opioid exposed placental immune cells towards regulatory and hyporesponsive states

We next performed DEG analysis between OUD_HCV+ and OUD_HCV-groups. The majority of DEGs were found among decidual macrophages, PAMM1B, and PAMM1A cells. First, upregulated genes with maternal HCV within the dMac_1 subset mapped to immune regulatory processes (*CD9, IL10, CCL20*) while downregulated genes mapped to immune effector processes, hemopoiesis and autophagy (*FCGR3A, C1QA, S100A8*) (Fig. S4A). Genes upregulated in the dMac_2 subset with HCV were important for responses to cytokine stimulus and inflammation, as well as cytokine production (*IL1RN, CXCL8, TNFAIP2*), while downregulated genes were associated with regulation of innate immune responses (*C1QA, HLA-C, CLEC12A*) (Fig. S4B). Finally, upregulated genes in the dMac_3 cluster were associated with clearance of apoptotic cells and debris, leukocyte cell-cell adhesion, and IL-6 production (*IL1RN, CXCL3, ICAM1*) in the OUD_HCV+ group (Fig. S4C). However, downregulated genes were associated with antigen processing/presentation, inflammatory/innate immune responses, and wounding (*HLA-DQA1, FCGR1A, S100A8*) (Fig. S4C). These findings suggest that overall decidual macrophage subsets are rewired towards a more regulatory phenotype with maternal OUD and HCV infection.

We next identified DEGs among PAMM subsets. Upregulated genes in the PAMM1B subset were associated with responses to cytokine and bacteria, as well as wound healing and chemotaxis (*IL-1, CXCL2, IL1RN*) in the OUD_HCV+ group (Fig. S4D). However, downregulated genes were associated with negative regulation of immune system processes, macrophage activation, and regulation of hemopoieses (*FOS, CEBPD, HLA-DQB1*) (Fig. S4D). Within the PAMM1A subset, upregulated genes mapped to positive regulation of cell migration, cytokine production, responses to hormones and type-II interferon (*EREG, CCL5, AREG*) in the OUD_HCV+ group (Fig. S4E). Downregulated genes in the PAMM1A subset with maternal OUD and HCV infection were associated with leukocyte activation, viral responses, and immune effector processes (*SPI1, HLA-DQA1, CYBA*) (Fig. S4E). These findings suggest that PAMM1B and PAMM1A shift towards a more activated phenotype with maternal OUD and HCV infection, characterized by high migratory and cytokine-producing capacity but suppressed antiviral functions.

In the T cell cluster, expression of genes associated with immune activation, including inflammatory responses, chemotaxis, and cell killing (*KLRK, CD8B, PIK3R1*) (Fig. S5A) were decreased, suggesting hyporesponsive T cells. In the dNK2 subset, expression of genes associated with antigen presentation, cell-cell adhesion, and regulation of viral entry into host cell was decreased (*GNLY, ID2, ITGAX*) in the OUD_HCV+ group (Fig. 5B), in line with dampened NK cell response to stimulation. Finally, we noted increased expression of genes associated with antigen presentation, antimicrobial responses and positive regulation of neutrophil migration (*HLA-DRB1, CXCL8, TNFAIP8*) with maternal OUD and HCV infection in the mDC subset (Fig. S5C). However, downregulated genes were associated with antigen processing/presentation, regulation of immune effector processes, and defense against bacteria (*ANXA2, FCGR1A, HLA-DRB5*) in the OUD_HCV+ group compared to OUD_HCV-(Fig. S4E-F). These findings indicate that mDCs are skewed towards supporting neutrophil trafficking while ability to activate T cells and mount an antimicrobial response is suppressed.

### Subhead 6: Vascular malperfusion and inflammation are abundant in placenta with maternal OUD±HCV exposure

Disruptions in placental structure have been shown to directly impact placental immunity (*46*). Therefore, we reviewed histopathological findings of decidua and surrounding chorionic villous tissues from healthy, term pregnancies (control) and those with OUD±HCV (Fig. 1A). Non-pathological tissues were more abundant in controls, as shown by healthy non-inflamed maternal decidua (Fig. 4A-B, arrows) and adjacent chorionic villous tissues exhibiting appropriate maturation and vascularity for gestation (Fig. 4B, box and arrows). An increase in pathological findings was associated with OUD±HCV (Fig. 4A), driven by fetal vascular malperfusion (FVM) including accelerated villous maturation and chorangiosis (Fig. 4C). Other notable findings include features of deciduitis/villitis (Fig. 4D, arrows) as well as maternal vascular malperfusion (MVM), including decidual arteriopathy of the basal plate and sites of increased perivillous fibrin (Fig. 4E, arrows) Overall, these findings suggest a detrimental impact on placental development with OUD±HCV, notably pathologies associated with placental hypoperfusion and hypoxia in the OUD_HCV+ group.

**Fig. 4:**
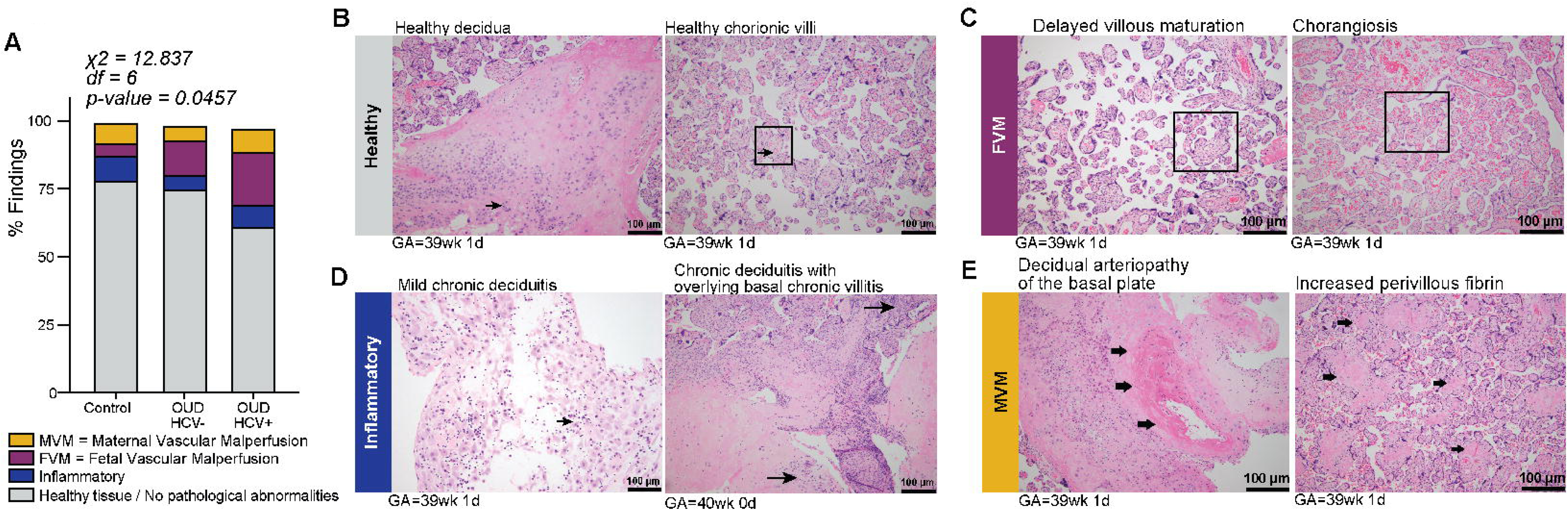
Maternal OUD±HCV is associated with inflammation and fetal/maternal vascular malperfusion. A) Stacked bar graph depicting the incidence of pathological findings. (B-E) Representative H&E staining of B) healthy, non-pathological villous tissues, C) features of fetal vascular malperfusion (FVM), D) inflammatory pathologies and E) features of maternal vascular malperfusion (MVM) identified in placental tissues from pregnancies with OUD±HCV.

### Subhead 7: The spatially resolved transcriptional landscape of placental cells reveals profound alterations associated with maternal OUD

Given the impact of maternal OUD±HCV on placental structure and the transcriptional profiles of placental immune cells, we assessed gene expression alterations in the context of placental architecture by Visium spatial transcriptomics (N=2 control, N=1 OUD_HCV-, N=1 OUD_HCV+) (Fig. 1A, Fig. S6A-B). Unique Manifold Approximation and Projection (UMAP) revealed 11 subpopulations (Fig. 5A, Fig. S6B, Table S2). Individual clusters were annotated using the EnrichR cell type database (Fig. S6C). We identified immune cells (leukocyte, myeloid, macrophage, and lymphoid), trophoblasts (cytotrophoblasts (CTB), syncytiotrophoblasts (STB), a combined STB/extravillous trophoblast (EVT) subset, as well as mature and immature EVT subsets), and structural cells (smooth muscle, endothelial) (Fig. S6C). Cluster frequencies were comparable between control and OUD groups, aside from a decrease in CTB abundance in the OUD group (Fig. 5B), a hallmark of placental dysfunction and stunted fetal growth (*47*).

**Fig. 5:**
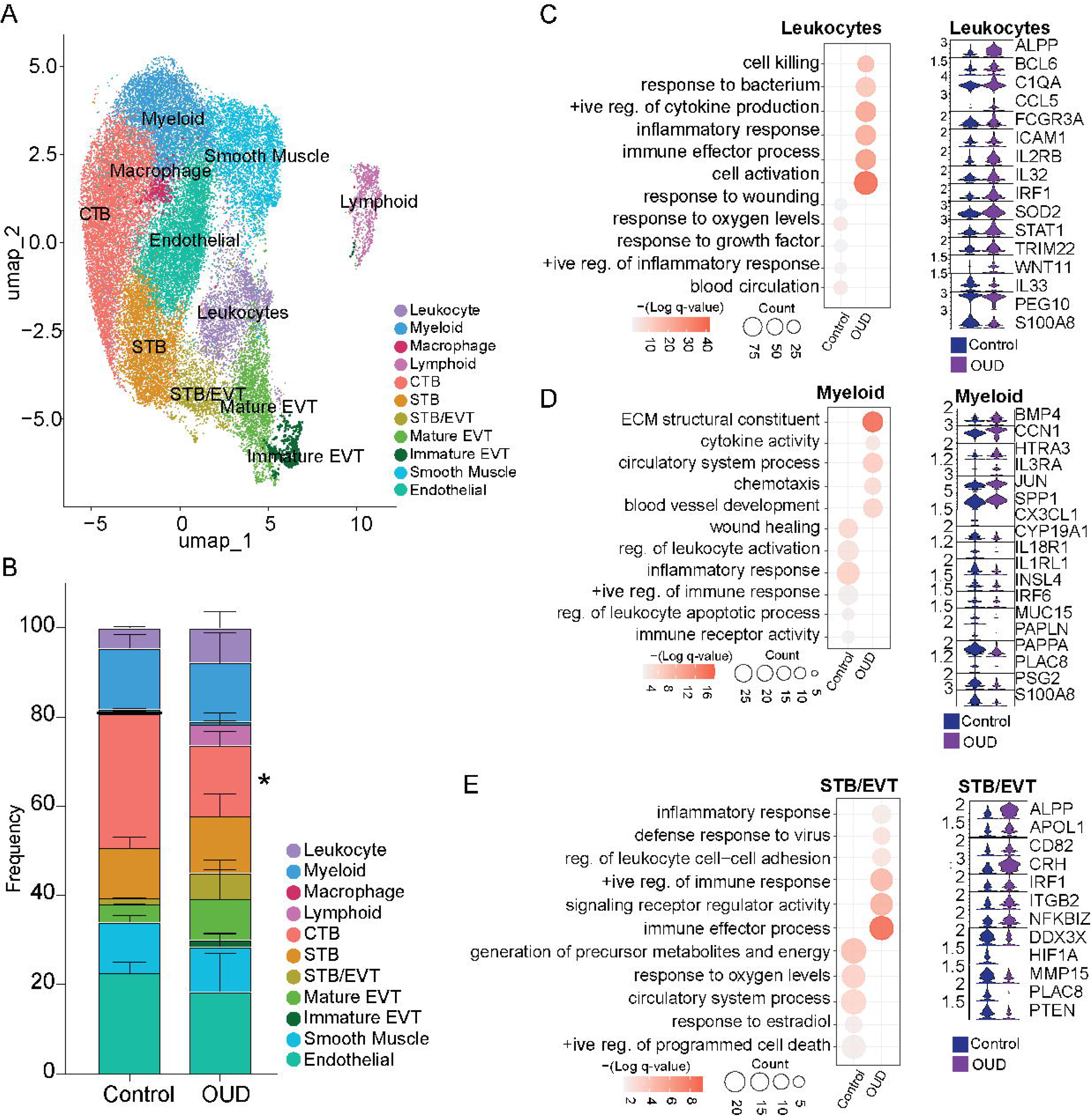
Maternal OUD alters the placental transcriptional profile on a spatial resolution. A) UMAP of identified clusters from placental tissue isolated from control and OUD samples (36,009 spots). B) Stacked bar graph of placental cluster frequencies between groups (control: 18,526 spots, OUD: 17,480 spots). (C-E) Bubble plots of DEGs between OUD and control groups (left) and violin plots showing representative DEGs from the select GO terms (right) for the shown cluster. For the bubble plots, the size of the bubble represents the number of genes associated with GO term and the color represents the significance (q-value).

DEGs upregulated in the Leukocyte cluster with maternal OUD enriched to GO terms associated with inflammation and immune activation, including cell killing and immune effector functions, bacterial responses, and cytokine production (*C1QA, ICAM1, FCGR3A*) (Fig. 5C). However, downregulated genes mapped to wounding and growth factors, oxygen-level responses, and regulation of inflammation (*IL33, PEG10, S100A8*) (Fig. 5C). Similarly, upregulated DEGs in the Lymphocyte subset played a role in cell activation and immune effector processes (*CCL3, GNLY, HAND2*), while downregulated genes were associated with vascular development and tissue remodeling (*VEGFC, FLT4, TRIM25*) (Fig. S7A) indicating a skewing towards an activated, pro-inflammatory effector state. These findings suggest that maternal OUD leads to heightened antimicrobial defenses and pro-inflammatory activities albeit reduced ability to mediate tissue repair. DEGs upregulated in the myeloid cluster with OUD were associated with extracellular matrix remodeling as well as cell migration, angiogenesis, and cytokine activity (*CCN1, IL3RA, SPP1*) (Fig. 5D). In contrast, genes important for immune responses and regulation of apoptosis were downregulated with maternal OUD, including *IRF8, PAPPA,* and *PSG2* (Fig. 5D), highlighting a shift among placental myeloid cells from immune defense and repair towards tissue remodeling, angiogenesis, and immune suppression.

Given that disrupted immune regulation by trophoblasts is associated with pregnancy complications, we next assessed DEGs among trophoblast clusters. In a healthy pregnancy, CTBs serve as a progenitor population that continuously differentiate to supply specialized trophoblasts to the syncytial surface and extravillous space. Here, upregulated genes in the CTB subset associated with differentiation, migration, and vascular remodeling (*CRH, FSTL3, GPER1*) but downregulated genes associated with NF-κβ activity, leukocyte chemotaxis, and inflammatory responses (*ERAP2, MAPK4, S100A8*) in the OUD group (Fig. S7B). With maternal OUD, upregulated genes in the STB cluster associated with syncytium formation, wound healing, and responses to estrogen and bacteria (*CRH, CYP19A1, FSTL3*) (Fig. S7C), highlighting that STBs in the OUD group are poised towards enhanced barrier function. We also identified a cluster containing both STBs and EVTs (STB/EVT). Our DEG analysis of this subset revealed upregulated genes associated with inflammatory response, antiviral defense, positive immune response, and immune effector processes with maternal OUD (*ALPP, APOL1, CD82*) (Fig. 5E). Genes downregulated with maternal OUD in the STB/EVT cluster were associated with metabolic pathways, response to oxygen and hormone levels, and apoptosis (*HIF1A, MMP15, PTEN*) (Fig. 5E).

Previous studies have characterized the differentiation of EVTs from proliferative, immature progenitors to mature, invasive cells that play critical roles in placental development (*48, 49*). In line with these studies, our analysis identified two EVT clusters delineated as immature and mature. While no DEGs were identified in the EVT_immature cluster, DEGs among the EVT_mature subset with maternal OUD include upregulated genes associated with heightened immune and inflammatory activity (*BCL6, CXCL12, MIF*) and downregulated genes associated with vascular remodeling and oxygen level responses (*CCR1, FOSB, IL33*) (Fig. S7D). These findings indicate that mature EVTs are poised towards heightened inflammatory signaling with reduced abilities for vascular remodeling and adaptation to low oxygen, potentially relevant to placental disorders marked by inflammation and insufficient trophoblast invasion (*50*).

Finally, placental structural cells support healthy placenta development as well as influence trophoblast and immune cell function. DEGs in the smooth muscle cell cluster, including increased expression of genes important for vascular remodeling and increased cytokine activity and chemotaxis (*PTGER1, TGFB3, SERPINE1*) and decreased expression of genes important for immune responses and wound healing (*KISS1, PEG10, CSHL1*) with maternal OUD compared to control (Fig. S7E). Similar trends were noted in the endothelial cell cluster, including increased expression of genes associated with blood vessel development, responses to hormones, and chemotaxis (*CD82, CXCL14, VCAM1*), while the expression of genes associated with responses to inflammation and pathogens, as well as cell proliferation and growth factor activity (*CD274, CTSH, GH2*) were decreased with maternal OUD compared to control (Fig. S7F). Overall, these findings suggest that OUD promotes vessel wall remodeling and inflammatory signaling with limited support for immune cell recruitment by smooth muscle and endothelial cell function favors. This phenotype is characteristic of pathological vascular remodeling, akin to atherosclerosis (*51*).

### Subhead 8: Maternal OUD disrupts cell communication networks in the term placenta

Given that cell signaling pathways were dysregulated with maternal OUD across cell types, we employed CellChat to infer changes in cell-to-cell communication from the Visium spatial transcriptomics data (*34*). Overall communication patterns across the 11 placental cell populations revealed increased total ligand-receptor interactions as well interaction strength in the OUD group, reflecting increased activation and cell recruitment (Fig. S8A). We next compared the outgoing and incoming signals of each subpopulation in control and OUD groups (Fig. 6A). The strength of total outgoing signals by all clusters was higher with maternal OUD (Fig. 6A), indicating a higher capacity to signals and shape the placenta microenvironment. Additionally, the strength of total incoming signals was higher across most immune cell clusters (leukocyte, lymphoid, macrophage), trophoblasts (EVT_mature, EVT_immature, STB_EVT), and structural cells (endothelial, smooth muscle) with maternal OUD (Fig. 6A). In contrast, the strength of total incoming signals to CTB and myeloid cell clusters was lower in the maternal OUD group (Fig. 6A). Collectively, these data indicate that maternal OUD globally intensifies placental intercellular communication potentially altering immune and trophoblast functions while dampening inputs to CTB and myeloid cells.

**Fig. 6:**
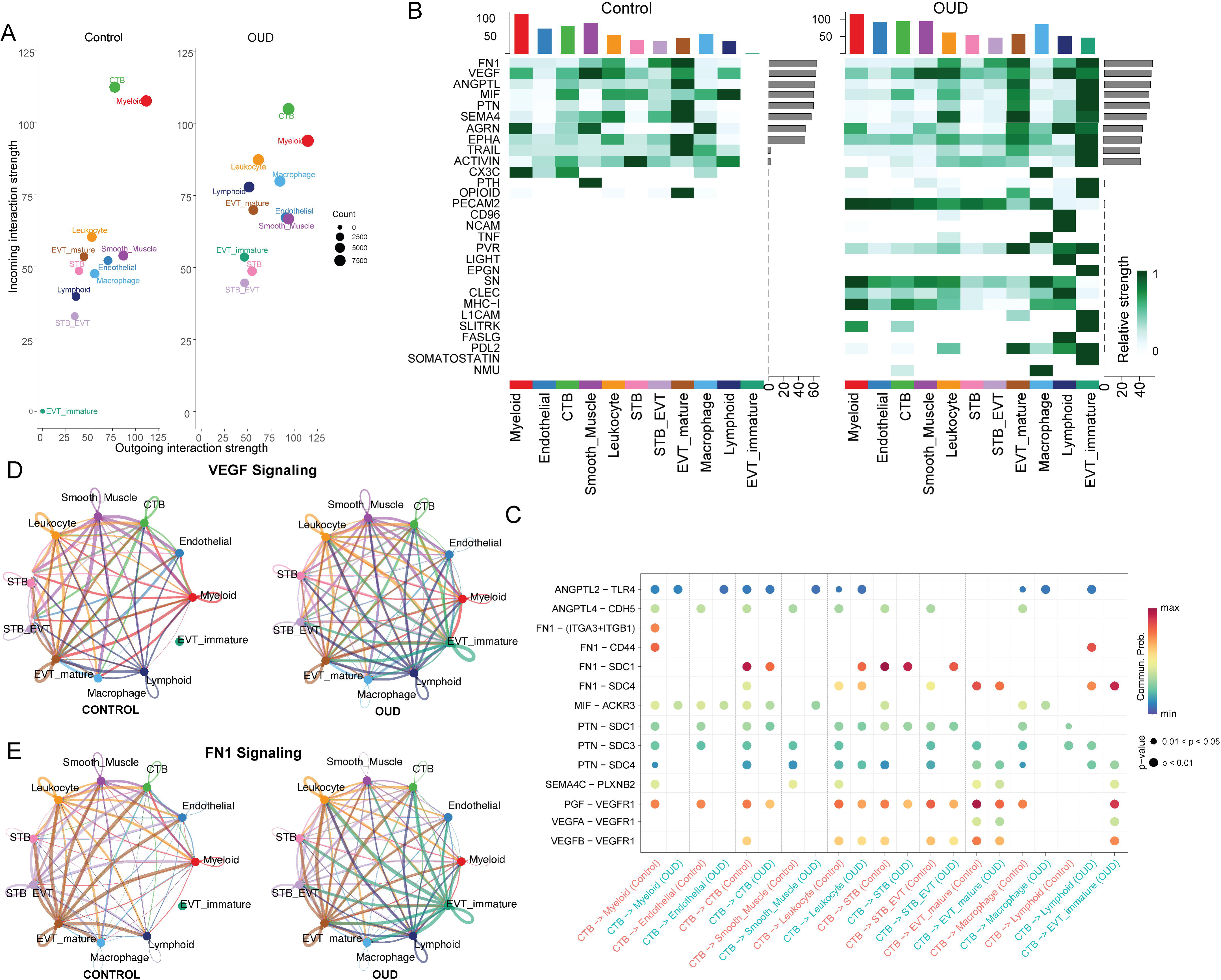
Maternal OUD impacts communication networks in the placenta. A) Scatterplot of the incoming and outgoing interaction strength of each cell population in control and OUD groups. The size of the node represents the total number of interactions in each type of cells. B) Heatmap displaying the overall signal strength of each signaling pathway in each cell population among control (left) and OUD (right) groups. Color bar plot (top) displays total signaling strength of specific cell clusters for all pathways. Gray bar plot (right) displays total signaling strength of specific pathways for all cell clusters. C) Scatter plot of the communication probability of select significant ligand-receptor pairs ligand-receptor pairs between control (left) and OUD (right) placenta, which contribute to the signaling from CTB to all other cell type clusters identified by our Visium spatial transcriptomics data. Dot color reflects communication probabilities and dot size represents computed p-values. Empty space indicates a communication probability of zero. (D-E) Circle plot of the inferred D) VEGF and E) FN1 signaling networks between cell types where the thickness of the line indicates greater number/communication probability and color of the line indicates origin of the signal.

We next compared the overall strength of signaling pathways in individual clusters between OUD and control groups (Fig. 6B). Interestingly, our analysis revealed more active overall signaling pathways (incoming and outgoing) in placentas exposed to maternal OUD, including PECAM2, PVR, SN, CLEC, and MHC-I. Signaling patterns important for trophoblast development and placental health across multiple cell clusters were stronger with maternal OUD, namely FN1, VEGF, MIF, and PDL2, among others (Fig. 6B). Furthermore, OPIOID signaling was lower in the EVT_mature subset but elevated in the EVT_immature subset with maternal OUD, suggesting that mature EVTs are possibly desensitized with opioid exposure (Fig. 6B). Interestingly, EVT_immature signaling pathways were strikingly enhanced in the maternal OUD group compared to control (Fig. 6B). All together, these findings suggest that overall placental immune-related signaling and pathways critical for trophoblast growth and vascular health are activated in the OUD group compared to control, where immature EVTs had the most pronounced signaling amplification underscoring altered development in early trophoblast stages.

We then identified significant ligand-receptor pairs that contribute to altered overall signaling pathways with maternal OUD (Fig. 6C, Fig. S8B). Our Visium spatial transcriptomics DEG analysis revealed altered expression of genes associated with cytokine receptor activity, cell activation, and growth factor signaling among trophoblast subsets with maternal OUD (Fig. 5E, Fig. S7). Overall signaling pathways showed a significant impact of maternal OUD on trophoblast signaling mechanisms, therefore, our downstream analysis focused on trophoblast cell clusters. Considering that CTBs are precursors to other, mature trophoblasts (STBs and EVTs), we first assessed differences in ligand-receptor signaling pathways within the CTB subset with maternal OUD (Fig. 6C). We identified FN1 and VEGF as a strong communication pathway between CTBs and EVT_immature cells with maternal OUD (Fig. 6B, D and E). FN1 and VEGF are both important regulators of trophoblast survival, migration and invasion, contribute to macrophage tissue repair functions, and are essential for placental vascularization and integrity (*52*). Therefore, we further explored the importance of these specific pathways in the context of maternal OUD. We show that CTBs also utilize FN1 and VEGF signaling pathways to interact with myeloid cells, endothelial cells, STBs, EVT_mature, and other CTBs in the control group, that are absent or dampened with maternal OUD (Fig. 6C, Fig. S8B), possibly as a result or cause of placental damage and inflammation. Interestingly, FN1 and VEGF signaling interactions were prominent in the endothelial cell cluster in the control group, but minimal in the OUD group (Fig. 6D and E, Fig. S8B). Overall, our CellChat analysis highlights that maternal OUD leads to aberrant placental signaling pathways indicative of disrupted trophoblast maturation and placental damage with maternal OUD.

## DISCUSSION

The human placenta is a maternal-fetal organ crucial for supplying nutrients to and removing waste from the developing fetus (*53*). Maternal OUD can negatively impact placental structure via direct and indirect mechanisms. Opioids have been shown to target endogenous opioid receptors on placental cells, including trophoblast (*54*) and immune cells (*55*), however the impact of these interactions on human placenta have not been well characterized (*56, 57*). Opioids can also indirectly impact placental function through disrupting the regulation of reproductive hormones, impairing placental blood flow, limiting trophoblast differentiation and invasion, and modulating inflammatory responses (*10*). Additionally, maternal HCV infection, commonly associated with intravenous drug use, can impair placental barrier (*58*). However, the impact of OUD on the maternal-fetal interface remains poorly understood.

Consistent with other reports of perinatal opioid (*59*) or HCV (*60*) exposure, newborns in our study were smaller, more likely to be admitted to intensive care units, suffer from neonatal opioid withdrawal syndrome (NOWS), require pharmacological interventions (morphine), and are diagnosed with congenital abnormalities at a higher rate. These adverse outcomes are likely mediated, in part, by abnormal placental development and therefore placental function throughout pregnancy (*61, 62*).

Our histological review per the Amsterdam criteria (*63*) revealed increased abundance of pathological findings in both OUD groups, notably features of FVM and villitis that is further exacerbated by HCV infection. Delayed villous maturation noted in opioid-exposed placenta was indicated by larger and less abundant terminal villi, restricting nutrient and oxygen exchange between mother and fetus (*64*). Fetal hypoxia is indicated by increased incidence of chorangiosis, a hallmark of low-grade hypoxia in placental tissue (*65*), as well as decreased expression of genes regulating oxygen-levels across leukocyte and trophoblast subsets with maternal OUD. Our scRNA-seq analysis revealed a depletion of CTBs which has been associated with placental hypoxia and reduced spiral artery remodeling, hallmarks of placental vascular malperfusion (*47*).

Furthermore, we show that CTB communication networks shift towards tissue remodeling and placental repair with maternal OUD, as shown by the stronger probability of PECAM2, SN, and FASLG signaling, among others, possibly contributing to poor maintenance of the placental barrier and immune function (*16*). These structural changes have significant implications for fetal health since delayed villous maturation has been recently identified as predictors of low scores on neurodevelopmental tests, intrauterine growth restriction (IUGR), and congenital abnormalities all of which are increased with maternal OUD (*66*).

To complement our histological findings, we interrogated the transcriptional landscape of the placenta on a special resolution. Structural cells showed increased expression of genes associated with wound healing and vasculature remodeling in non-immune cells of the placenta with maternal OUD. These changes could be underlying vascular malperfusion prevalent in this cohort. Indeed, chronic inflammation is strongly associated with FVM and vessel wall thickening (*67*). In addition to maintaining the placental barrier, structural cells of the placenta closely interact with surrounding trophoblasts and immune cells to mediate pathogen responses, control inflammation, and maintain fetal tolerance (*68*). However, gene expression signatures across smooth muscle and endothelial cell clusters suggests that OUD disrupts immune cell recruitment and antimicrobial responses. Furthermore, placental endothelial and trophoblast cells have reduced FN1 signaling to CD44 expressed by placental myeloid cells with maternal OUD. The FN1-CD44 pathway links regulatory macrophage function with trophoblast survival and placental architecture, providing a possible mechanism underlying increased placental pathologies with maternal OUD.

Additionally, CTBs are precursors to EVTs that penetrate the decidua and remodel spiral arteries to ensure adequate blood supply to the developing fetus, as well as STBs that form the outer layer of placental villi and are especially adapted to shield the fetus from the maternal immune system (*67*). Stronger outgoing signals from CTBs to immature EVTs were prominent in the OUD group, particularly those involved in tissue repair pathways. Notably, VEGFB and PlGF signals bind exclusively to VEGFR1 and excessive signaling along these pathways have been associated with aberrant angiogenesis and enlarged maternal vascular spaces (*69*), in line with our findings of increased incidence of MVM with maternal OUD. In addition, transcriptional profiles of STBs were indicative of barrier dysfunction and wound healing while the transcriptional signatures of EVT_mature suggested heightened immune and inflammatory signaling but reduced vascular remodeling processes, akin to insufficient trophoblast invasion (*70*). Insufficient trophoblast differentiation/function is prominent placental pathologies that have been associated with opioid use during pregnancy, including fetal growth restriction, placental abruption, and preterm birth (*48*). Therefore, our observations indicate that maternal OUD and HCV infection results in the rewiring of CTBs which in turn disrupts the function of EVT/STBs leading to vascular malperfusion, placental hypoxia, and fetal growth restriction (*71*).

Various cytokines produced by placental cells can influence the inflammatory milieu, immune responses, and placental barrier function (*72*). We report elevated concentrations of pro-inflammatory factors with maternal OUD in the decidua. While a pro-inflammatory milieu is important for initiating labor and parturition in late pregnancy, exacerbated markers of inflammation are noted in several complications including pre-eclampsia and recurrent miscarriage (*73*). Notably, in vitro models using first trimester placenta showed that elevated decidual TNF impairs trophoblast recruitment and function as well as the expression of angiogenesis factors (*74*).

Dysregulated cytokine secretion with maternal OUD is less profound in the (villous) fetal compartment. Low concentrations of IL-6 and IL-8 impede trophoblast migration and angiogenesis and have been associated with impaired fetal development and immune-endocrine crosstalk (*75*). On the other hand, elevated levels of IL-12 and IL-7 have been associated with fetal growth restriction (*76*), in line with small for gestational age newborns and vascular malperfusion with maternal OUD. Furthermore, these findings align with dysregulated expression of genes important for repairing tissue damage and responding to hypoxia/growth factors across most cell subsets identified by our spatial transcriptomics analysis.

Among immune cell populations, we report a decreased frequency of the regulatory, dMac_3 macrophage subset with maternal OUD and HCV infection by flow cytometry. Given that regulatory macrophages suppress inflammation and support vascular growth, the depletion of dMac_3 cells with maternal OUD could explain elevated concentrations of inflammatory markers and impaired tissue-repair functions. Moreover, loss of regulatory macrophages has been linked to placental insufficiency (*77*). These changes in immune cell frequency detected by scRNA-seq with maternal OUD±HCV were not as prominent as those measured by flow cytometry. This discrepancy may be attributed to the fact that protein markers can remain stable despite fluctuations in transcriptional profiles (*78*). Nevertheless, single-cell RNAseq analysis of decidual macrophage subsets showed upregulation of processes associated with immune activation and inflammatory responses across all subsets with maternal OUD. However, antimicrobial responses by decidual macrophages to *ex vivo* bacterial TLR stimulation were dampened with maternal OUD±HCV infection, suggesting an immune tolerant-like state. Activated but functionally defective decidual macrophages have been associated with multiple placental pathologies, including villitis, vascular malperfusion, disrupted cytokine/chemokine signaling networks, and trophoblast differentiation (*79*), all of which align with our findings with maternal OUD.

Increased monocyte recruitment to the placenta has been associated with impaired placental perfusion, fibroblast proliferation, chronic inflammation, and hypoxia (*80*). PAMM1B cell transcriptional profiles highlight increased cytokine signaling with maternal OUD and HCV infection indicating a shift towards differentiation to PAMM1A cells. Furthermore, transcriptional profiles of PAMM1B, PAMM1A, as well as PAMM2 cells had heightened tissue repair and placental development pathways. Maternal monocytes and macrophages contribute to maintaining immune tolerance, pathogen defenses, and labor onset through interactions with decidual T cells (*81*). We report dampened CD4 and CD8 T cell functional responses to stimulation with maternal OUD±HCV, suggesting an immune tolerant or exhausted T cell phenotype. This was further reflected by our scRNAseq data where expression of genes important for immune responses and cell killing were dampened with maternal OUD. Additionally, our findings indicate decreased abundance of CD56^+^CD16^+^ dNK cells, a unique dNK cell subset that are poor cytokine producers but are highly cytotoxic, important for responding to infection, and regulating trophoblast invasion (*45*). Here, our data showed decreased abundance and poor degranulation capacity of CD56^+^CD16^+^ dNK cells with maternal OUD±HCV. In line with these findings, our scRNAseq data showed that the cytotoxic, dNK2 subset had lower expression of genes associated with migration and antimicrobial responses, consistent with poor cytotoxic responses among decidual NK cells.

In summary, this study used multi-omic approaches to uncover the impact of OUD±HCV on placental structure and immune function. Our findings shed light on the complex interplay between endothelial, smooth muscle, trophoblast, and immune cells, providing a framework for possible mechanisms underlying poor placental development with maternal inflammatory conditions. Maternal OUD, particularly when combined with HCV infection, profoundly alters the immune and structural landscape of the placenta, driving adverse pregnancy and neonatal outcomes through chronic inflammation, immune dysregulation, and impaired placental vascularization. While we were able to control for many maternal factors including maternal age, mode of delivery, fetal sex, pre-pregnancy BMI, and maternal HCV infection, we were unable to stratify by additional maternal factors, without compromising statistical power. For instance, parity was significantly higher in both OUD groups compared to control. Additionally, polysubstance use is prevalent with people who use drugs, including high rates of THC, nicotine, alcohol, and stimulant use. Future studies in animal models should be explored to better control for these confounding variables.

## MATERIALS AND METHODS

### Sample collection

Placental biopsies were collected from full term pregnancies. Leukocytes from decidua basalis and chorionic villous tissues were isolated using previously established methods (*21, 82*). Additional segments of placental tissue were collected, and flash frozen for downstream tissue homogenization. A third piece of placental tissue was placed in cassettes for fixation in 10% formalin. Fixed tissues were then paraffin embedded for subsequent H&E staining. PBMCs and plasma were isolated from maternal blood samples collected at delivery as previously described (*21*).

### Assessment of placental structure and histology

Placental histology was assessed by review of formalin fixed, paraffin embedded, and H&E-stained placental tissues. Placental lesions were documented and classified per the Amsterdam Consensus Statement Guidelines (*63*).

### Luminex assays

Supernatants of homogenized flash frozen placental tissues were collected, as previously described (*43*), for Luminex assay per manufacturer’s instructions: R&D Human Luminex® Discovery Assay, Cat#:LXSAH (inflammatory) and R&D Human Luminex® Discovery Assay, Cat#:LXSAHM (angiogenesis). Markers of placental function/development in maternal circulation were measured in maternal plasma using the following assays per manufacturer’s instruction: R&D Human Luminex® Discovery Assay, Cat#:LXSAHM (angiogenesis), PAPP-A Human ProcartaPlex™ Simplex Kit, Cat#:EPX010-12393-901, and Millipore Human Angiogenesis and Growth Factor Panel, Cat# HAGP1MAG-12K. sPLSDA from Luminex data was performed using the MixOmics (*83*) package with the normalized counts.

### Phenotyping by flow cytometry

Placenta leukocytes were thoroughly washed with FACS buffer and stained with a cocktail containing the following antibodies at a ratio of 1:20: Decidua: CD45, CD4, CD8b, CD14, HLA-DR, CD11c, CD56, CD16, FOLR2, S100A8/9, and CD9; Chorionic villous: CD45, CD14, HLA-DR, FOLR2, CD9, and CCR2 (BioLegend). True-Stain Monocyte Block and Human TruStain FcX™ (BioLegend) were also added to the surface staining cocktail (1:20). After incubation for 20min at 4°C, cell pellets were washed with FACS buffer and run using an Attune NxT and analyzed on FlowJo 10.10 (BecktonDickinson).

### Ex vivo stimulation and intracellular cytokine staining

1×106 decidua or chorionic villous leukocytes were stimulated for 16h at 37°C in 10% FBS:RPMI with or without bacterial TLR cocktail containing 1mg/mL LPS-B5 (TLR4, cat#:tlrl-b5lps, Invivogen), 2mg/mL Pam3CSK4 (TLR1/2, cat#:TLRL-PMS, Invivogen), and 1mg/mL FSL-1 (TLR2/6, cat#:SML1420, Sigma). Brefeldin A was added after 1h incubation and cells were cultured for an additional 15h at 37°C before surface and intracellular staining. Decidual leukocytes were stained with the following surface antibodies (1:20): CD45, CD2, CD20, CD14, HLA-DR, CD11c, CD9, and CCR2 (BioLegend) for 30min in the dark at 4°C. Chorionic villous leukocytes were stained with the following surface antibodies (1:20): CD45, CD14, HLA-DR, FOLR2, CD9 and CCR2 (BioLegend) for 30min in the dark at 4°C. Samples were fixed and permeabilized using fixation and permeabilization wash buffer (BioLegend) at 4°C for 20min and stained intracellularly for 4h for TNFα and IL-6 (BioLegend)(1:20). Samples were then washed, acquired, and analyzed as outlined for phenotyping.

1×106 decidual leukocytes were stimulated for 16h at 37°C in 10% FBS:RPMI in the presence or absence of PMA/ionomycin. Brefeldin A was added after 1h incubation and cells cultured for an additional 15h at 37°C. To delineate decidual leukocytes, cells were stained with the following surface antibodies: CD45, CD2, CD20, CD14, HLA-DR, CD56, CD107a, CD4, and CD8 (BioLegend)(1:20) for 30min at 4°C. Samples were fixed, permeabilized, and stained intracellularly for IL-2, IL-17, TNFα, and IFNγ (1:20) in permeabilization buffer for 4h at 4°C. Cells were then washed, acquired, and analyzed as outlined for phenotyping.

### Phagocytosis Assay

Chorionic villous leukocytes were incubated for 2h at 37°C in media containing 1 mg/mL pH-sensitive pHrodo E.Coli BioParticles conjugates (ThermoFisher). Cells were stained with antibodies against CD14, HLA-DR, FOLR2, CD9, and CCR2 (BioLegend)(1:20), then resuspended in ice cold FACS buffer. Cells were then washed, acquired, and analyzed as outlined for phenotyping.

### CD45 3’ Single Cell RNA library preparation

Placental immune cells were thawed, stained with CD45-FITC at 4°C in 1% FBS:DPBS and sorted into 30% FBS:RPMI using the BD FACS Aria-3 cell sorter. Cells were counted in triplicate and an equivalent number of cells were pooled by group (control and OUD) and resuspended in 0.4% BSA:PBS in a final concentration of 1200 cells/μL, and immediately loaded on the 10x Genomics Chromium Controller with a loading target of 30,000 cells. Libraries were generated per the manufacturer’s instructions (V3.1, 10x Genomics). Libraries were sequenced using the Illumina NovaSeq6000 with a sequencing target of 20,000 reads/cell.

### Single Cell RNA sequencing analysis

Raw reads were aligned and quantified using the CellRanger Software Suite (V6.0.1, 10X Genomics) against the GRCh38 human reference genome. Downstream processing of aligned reads and QC was performed using Seurat (V5.1.0) as previously described (*21*). Data objects for the placenta were integrated to remove leukocytes form infiltrating peripheral blood as previously described (*21*). Data normalization, dimensional reduction, cell type assignment (Table S1) and differential gene expression analysis (Table S1), were performed as previously described (*21*).

### Visium spatial transcriptomics library preparation

Placental architecture was assessed by Visium spatial transcriptomics (10x Genomics) with CytAssist using Demonstrated Protocol CG000520. RNA was extracted using the Qiagen FFPE RNA kit per manufacturers protocol. Blocks with a DV200 >40% were deparaffinized and H&E stained. Imaging was performed on a Nikon Ni-E microscope and image tiles stitched with Nikon Elements software. Slides were de-crosslinked and immediately hybridized with the Visium Human Transcriptome Probe Kit V2 (PN-1000466, 10x Genomics) per manufacturer instructions. Libraries were sequenced using the Illumina NovaSeq6000 with a sequencing target of 25,000 paired reads per covered spot.

### Visium spatial transcriptomics analysis

Raw reads were aligned and quantified using the SpaceRanger Software Suite (V3.1.1, 10X Genomics) against the GRCh38 human reference genome. Downstream processing of aligned reads and QC was performed using Seurat (V5.1.0) as outlined for scRNA-seq above and as previously described (*21*). Data objects for the placenta were integrated to remove leukocytes form infiltrating peripheral blood as previously described (*21*). Data normalization, dimensional reduction, cell type assignment (Table S2) and differential gene expression analysis (Table S2), were also performed as previously described (*21*).

### Cell-Cell Interaction Analysis

CellChat ^19^ was employed to infer probabilities of intercellular communication between clusters identified by spatial transcriptomics. A CellChat ^19^ object was generated from a Seurat V5 object with the createCellChat function. Data was preprocessed with the function identifyOverExpressedGenes and identifyOverExpressedInteractions and communication probabilities between clusters were determined with computeCommunProb (truncatedMean, trim=0.1, interaction, range=250, contact.range=100). Communications were filtered to a minimum number of 10 cells and signaling pathway probabilities were calculated. Data was aggregated for interaction comparisons between groups (Control and OUD).

## LIST OF SUPPLEMENTARY MATERIALS

**Table S1:** Marker genes and DEG lists supporting the single-cell RNA sequencing data in Fig. 3-4 and Fig. S3-4.

**Table S2:** Maker genes and DEG lists supporting the Visium spatial transcriptomics data in Fig. 6 and Fig. S5.

## ACKNOWLEDGMENTS

We are grateful to all participants in this study. We thank the MFM Research Unit at University of Kentucky for sample collection and members of the Messaoudi Laboratory for assistance with tissue processing. Additionally, this research was supported by the University of Kentucky Biospecimen Procurement & Translational Pathology Shared Resource Facility of the Markey Cancer Center (P30CA177558), the Arts & Sciences Imaging Center, as well as the Center for Computational Sciences and Information Technology Services Research Computing Morgan Compute Cluster and associated research computing resources.

## Author contributions

Conceptualization, I.M, J.O.B, C.C.; methodology, I.M, J.O.B, C.C.; investigation, H.E.T., B.M.D, S.W., N.R.S, M.E.B.; writing H.E.T., B.M.D, S.W., D.C.M., N.R.S., and I.M.; funding acquisition, H.E.T. and I.M.; participant enrollment, C.C. and J.O.B. All authors have read and approved the final draft of the manuscript.

## Data availability

The datasets supporting the conclusions of this article are available on NCBI’s Sequence Read Archive: single-cell RNA sequencing (PRJNA1234184), Visium spatial transcriptomics (PRJNA1231418).

## Conflict of interest

The authors declare that the research was conducted in the absence of any commercial or financial relationships that could be constructed as a potential conflict of interest.

## Funding

This study was supported by grants from the National Institutes of Health: 1R01DA059152-01 (IM), 7R01AI145910-05S1(IM), TL1TR001997 (HT) and pilot funding from the University of Kentucky, including the Clinical and Translational Science Substance Use Disorder pilot grant 3210003238 (IM). This research was indirectly supported by the Kentucky Opioid Response Effort (KORE) via Substance Abuse and Mental Health Services Administration (SAMHSA) Grants, H79TI081704, H79TI083283, as well as the data management system that is hosted by UK with grant support from NIH CTSA UL1TR001998. The content is solely the responsibility of the authors and does not necessarily represent the official views of the National Institutes of Health or the University of Kentucky.

## STATISTICAL ANALYSES

Comparisons were made between controls and pregnancies with OUD±HCV infection. Normality was assessed using the Shapiro-Wilk test (alpha=0.05) and outliers identified through ROUT analysis (Q=0.1%). If the dataset was normally distributed, group differences were assessed using an unpaired T-test with Welch’s correction (2-groups) or one-way ANOVA (3-groups). In cases where the data did not meet Gaussian assumptions, group comparisons were conducted using the Mann-Whitney (2-groups) or Kruskall-Wallis (3-groups). Group comparisons based on categorical data were performed via Chi-square tests. The threshold for statistical significance was p<0.05 and comparisons were considered trending when p<0.1. Statistical analyses were performed using Prism (GraphPad).

---

**Fig. S1:**
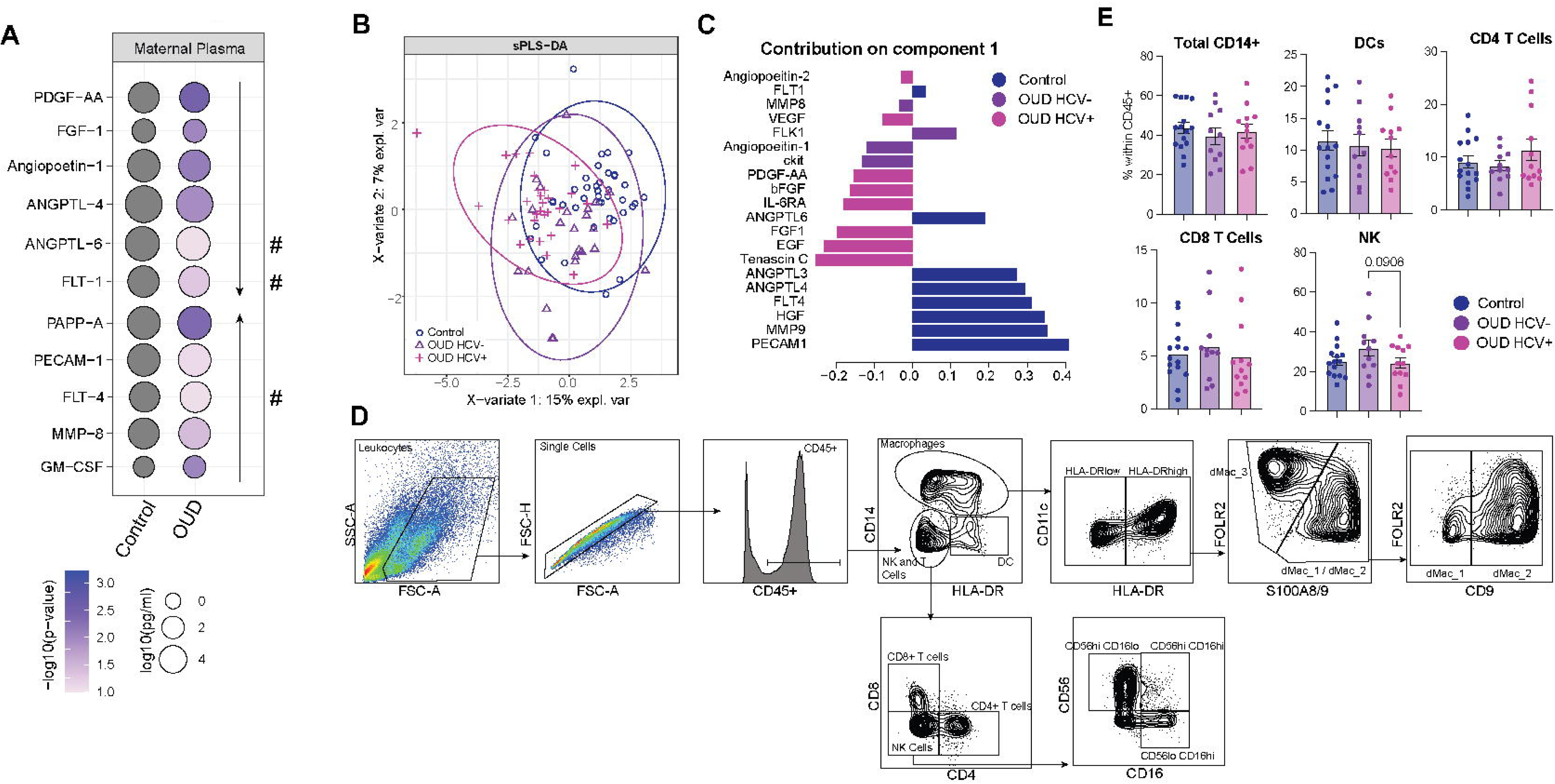
Phenotype and function of decidual leukocytes are altered by maternal OUD±HCV. A) Bubble plot of markers of placental dysfunction detected in maternal plasma between control and OUD groups. The size of the bubble represents the concentration (pg/ml) where the color represents the significance (p-value). B) PCA from the sPLSDA comparisons of control, OUD_HCV-, and OUD_HCV+ groups by concentration of markers of placental dysfunction measured in maternal plasma by luminex assay displayed in 1-2 space. C) Bar plot of markers of placental dysfunction delineated by group from the sPLS-DA component 1. D) Representative gating strategy for the identification of decidual immune cell subsets by flow cytometry. E) Bar plots of the frequency of total CD14+, dendritic cells (DC), CD4+ T cells, CD8+ T cells, and NK cell subsets within the CD45+ population.

**Fig. S2:**
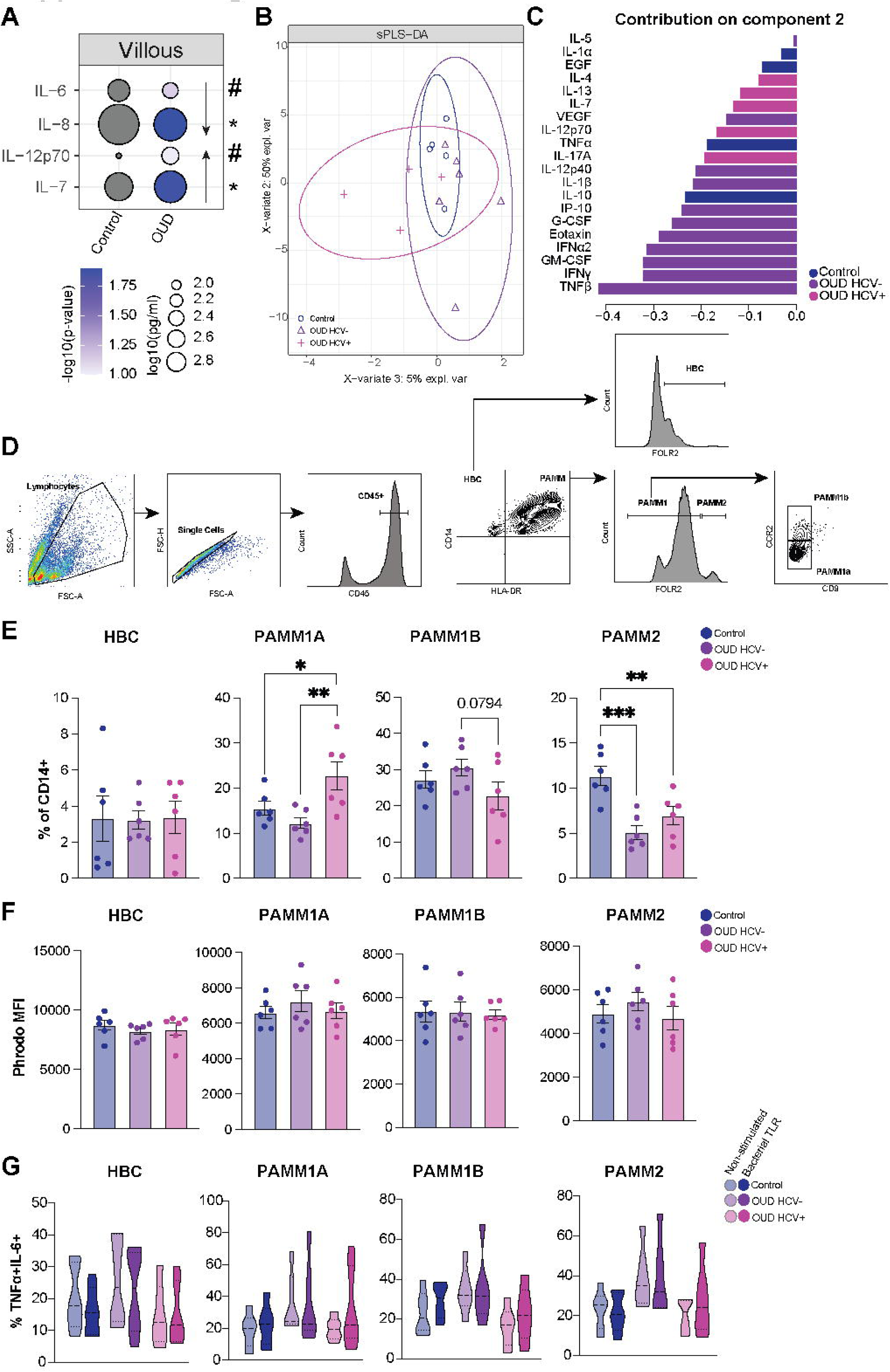
Immune phenotype and function of villous leukocytes are altered by maternal OUD±HCV. A) Bubble plot of immune mediators detected in villous tissue homogenate supernatant. The size of the bubble represents the concentration (pg/ml) where the color represents the significance (p-value). B) PCA from the sPLSDA comparisons of markers of placental dysfunction measured in placental tissue homogenate by luminex assay displayed in 1-2 space. C) Bar plot of markers delineated by group from the sPLS-DA component 2. D) Representative gating strategy for the identification of villous immune cell subsets by flow cytometry. E) Bar plots of CD14+ villous immune cell subset frequencies. F) Bar plots of PhRodo expression (MFI) at resting among chorionic villous immune cell subsets. G) Violin plots depicting lack of total TNFα and IL-6 responses to bacterial TLR stimulation among chorionic villous immune cell subsets with maternal OUD±HCV and in controls. ^#^=p<0.1, *=p<0.05, **=p<0.01, ***=p<0.001

**Fig. S3:**
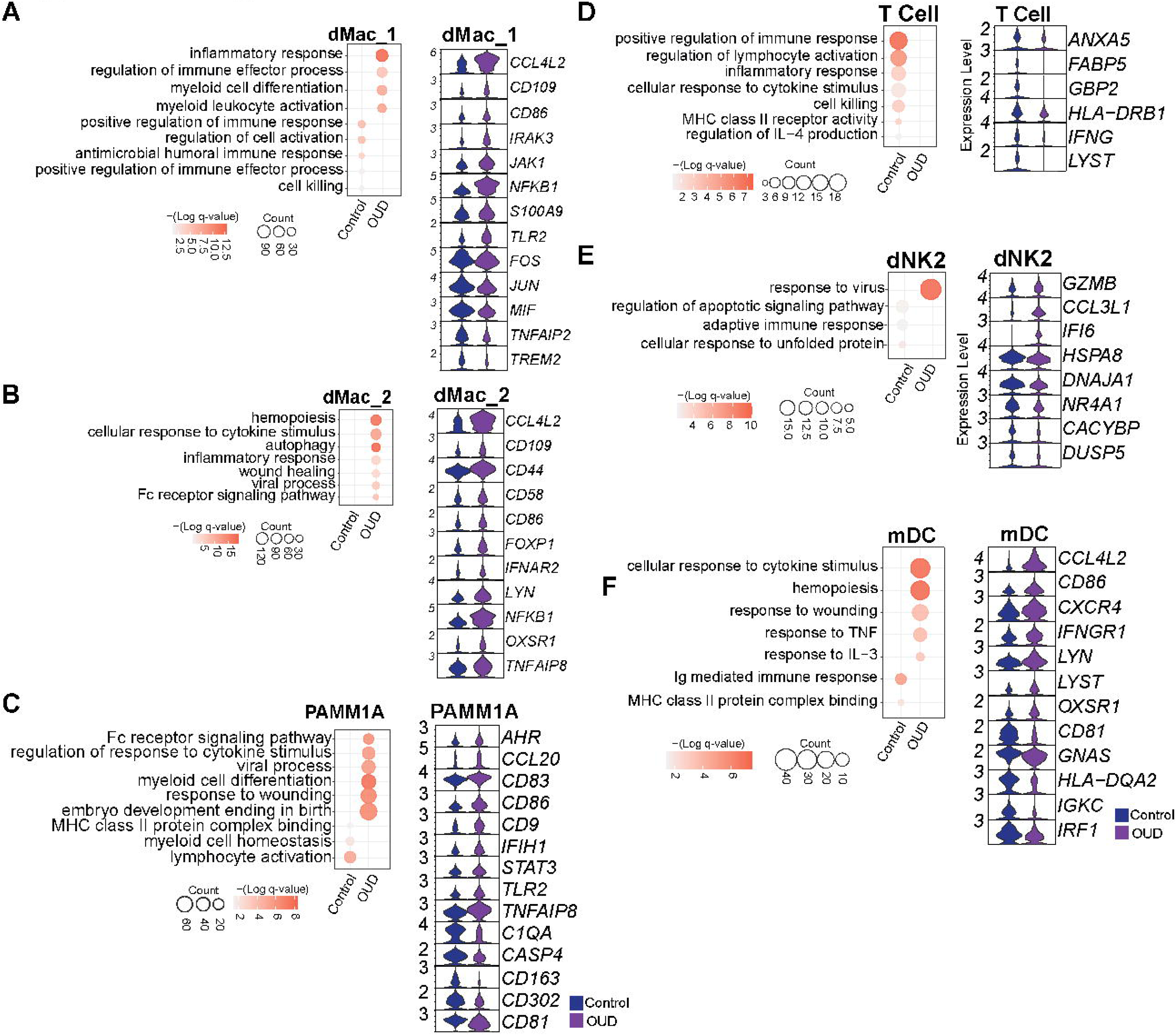
Maternal OUD alters the placental immune landscape. (A-F) Bubble plots of DEGs between control and OUD groups for select clusters (left) and violin plots showing representative DEGs from the select GO terms for the shown cluster (right). For the bubble plots, the size of the bubble represents the number of genes associated with GO term and the color represents the significance (q-value).

**Fig. S4:**
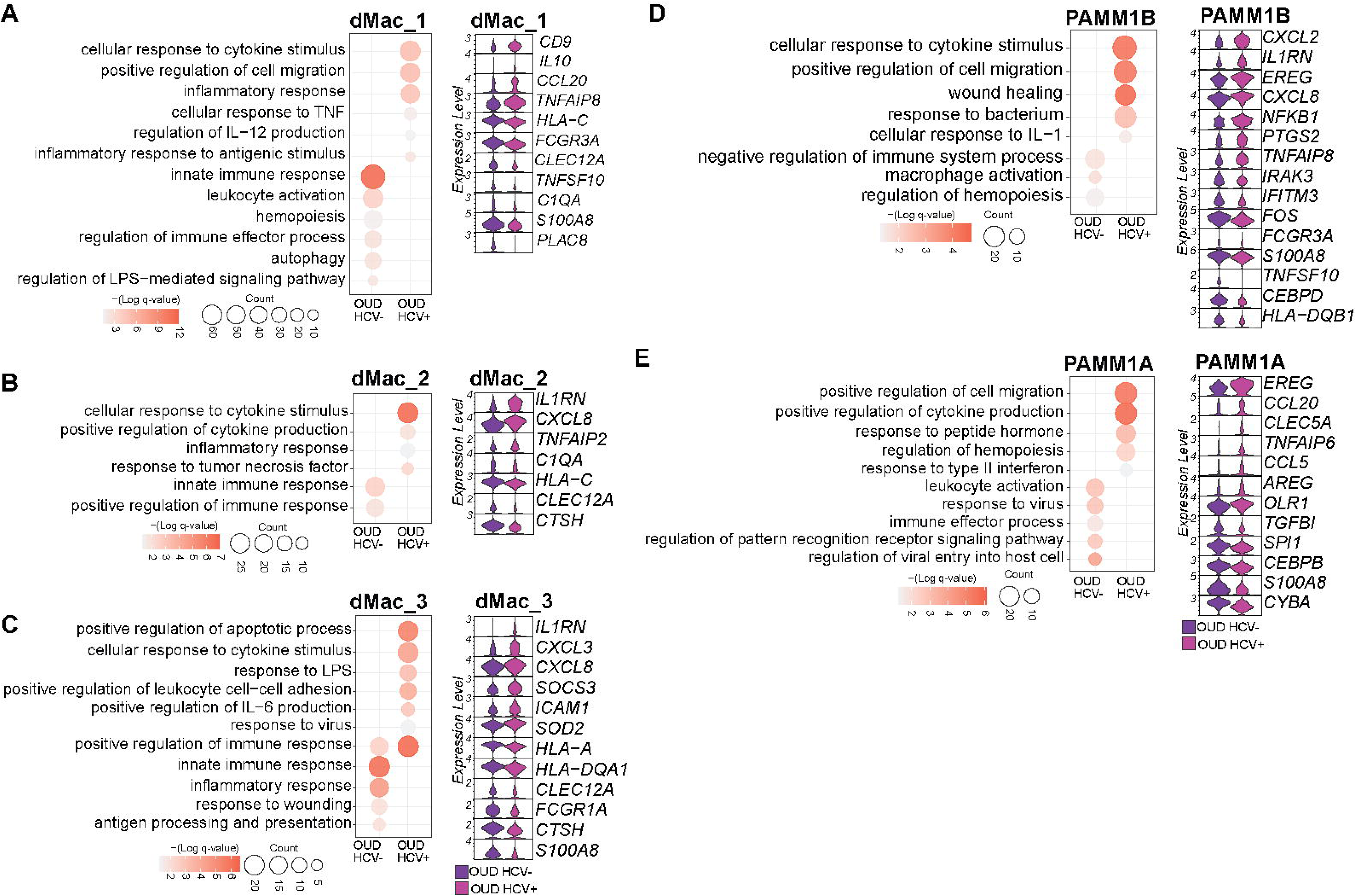
Concurrent maternal HCV infection impacts the transcriptional profile of innate placental immune cells. (A-E) Bubble plots of differentially expressed genes (DEGs) between OUD_HCV- and OUD_HCV+ groups (left) and violin plots showing representative DEGs from the select GO terms (right) for the shown cluster. For the bubble plots, the size of the bubble represents the number of genes associated with GO term and the color represents the significance (q-value).

**Fig. S5:**
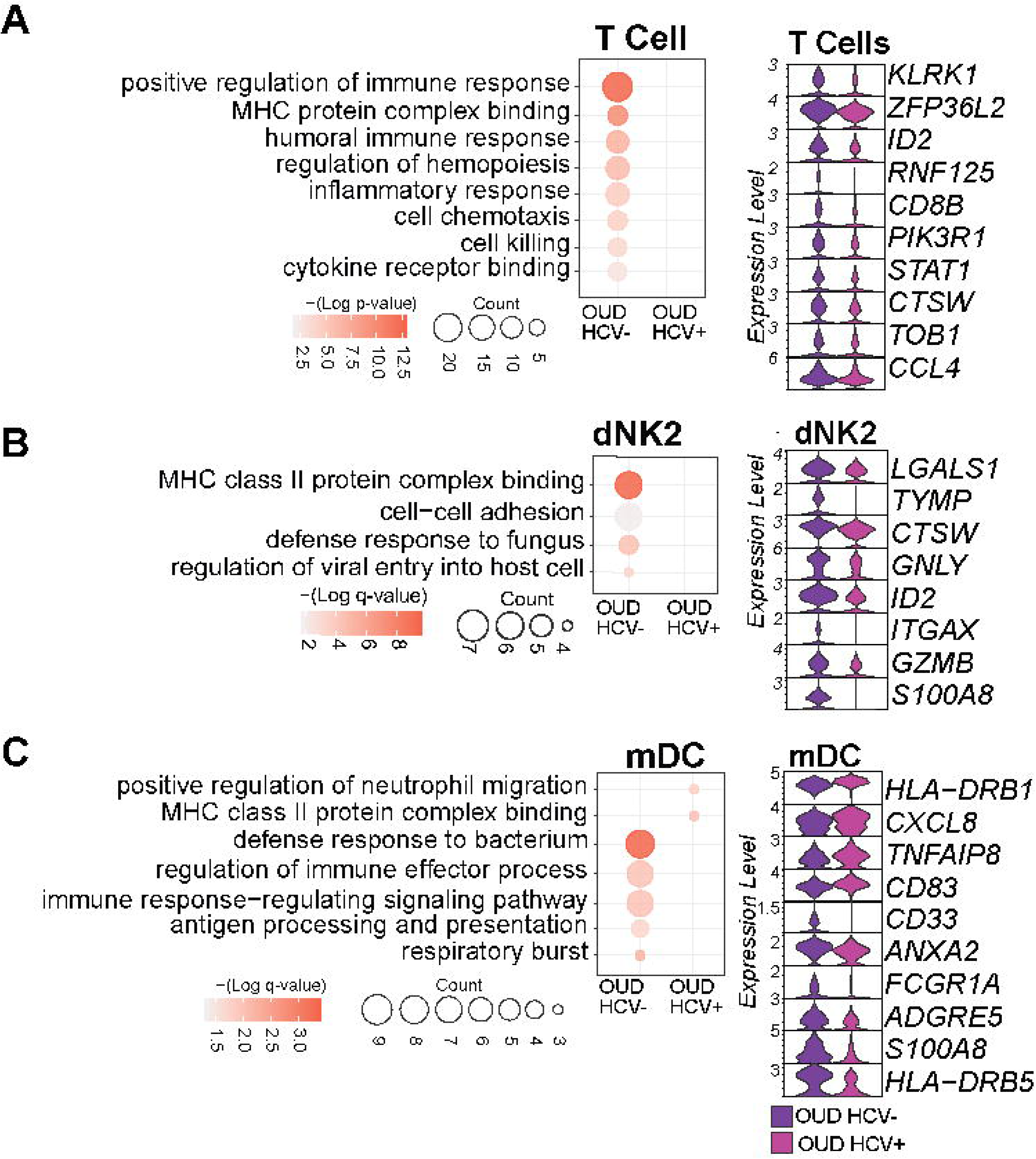
Concurrent maternal HCV infection impacts the transcriptional profile of adaptive placental immune cells. (A-C) Bubble plots of DEGs between OUD_HCV- and OUD_HCV+ groups (left) and violin plots showing representative DEGs from the select GO terms (right) for the shown cluster. For the bubble plots, the size of the bubble represents the number of genes associated with GO term and the color represents the significance (q-value).

**Fig. S6:**
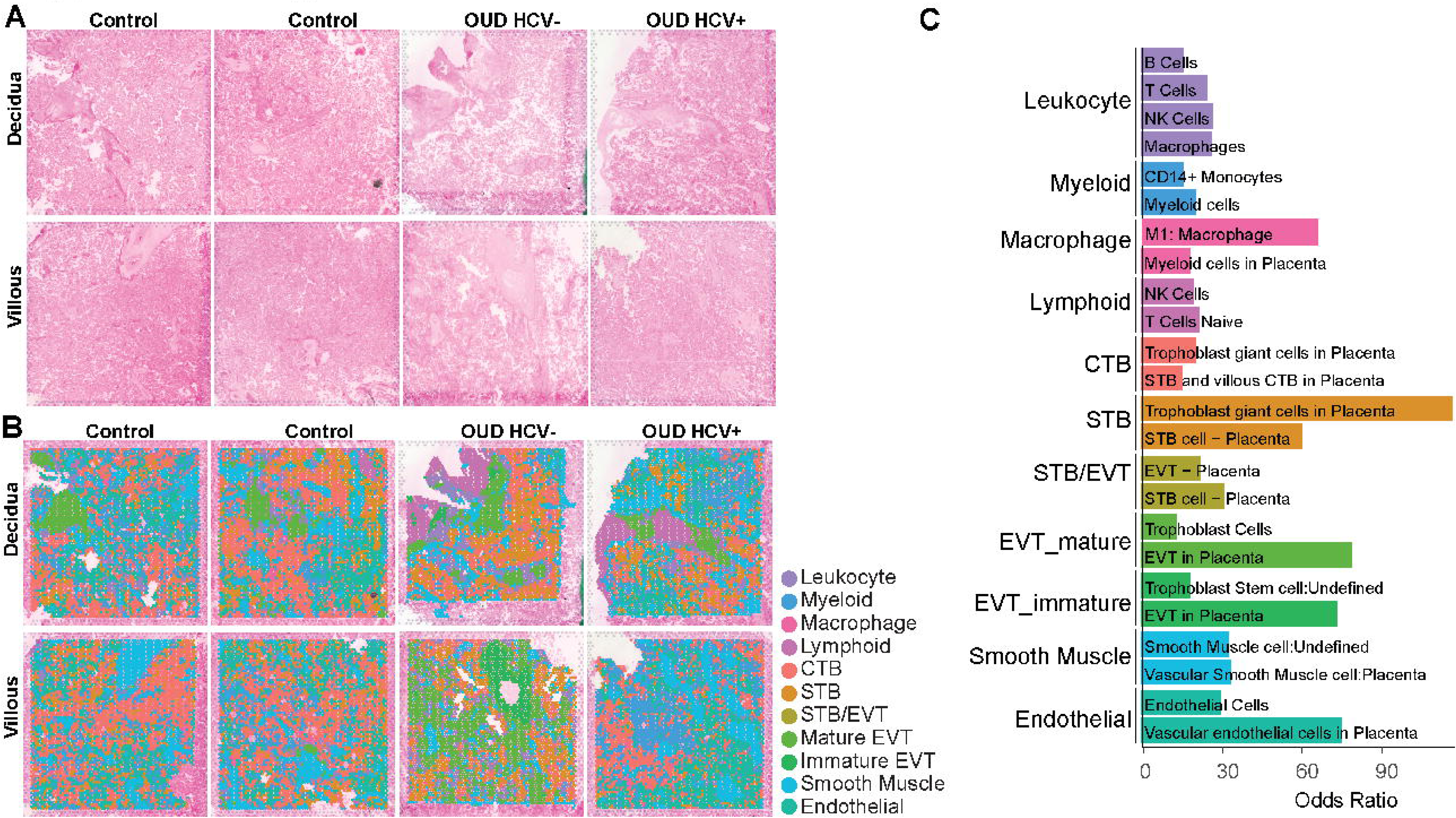
The placental transcriptional landscape at a spatial resolution. A) H&E-stained sections of placental tissues. B) H&E-stained sections overlayed with cluster identity spots. C) Bar plot of EnrichR cell type database odds ratios for marker genes from the indicated cluster.

**Fig. S7:**
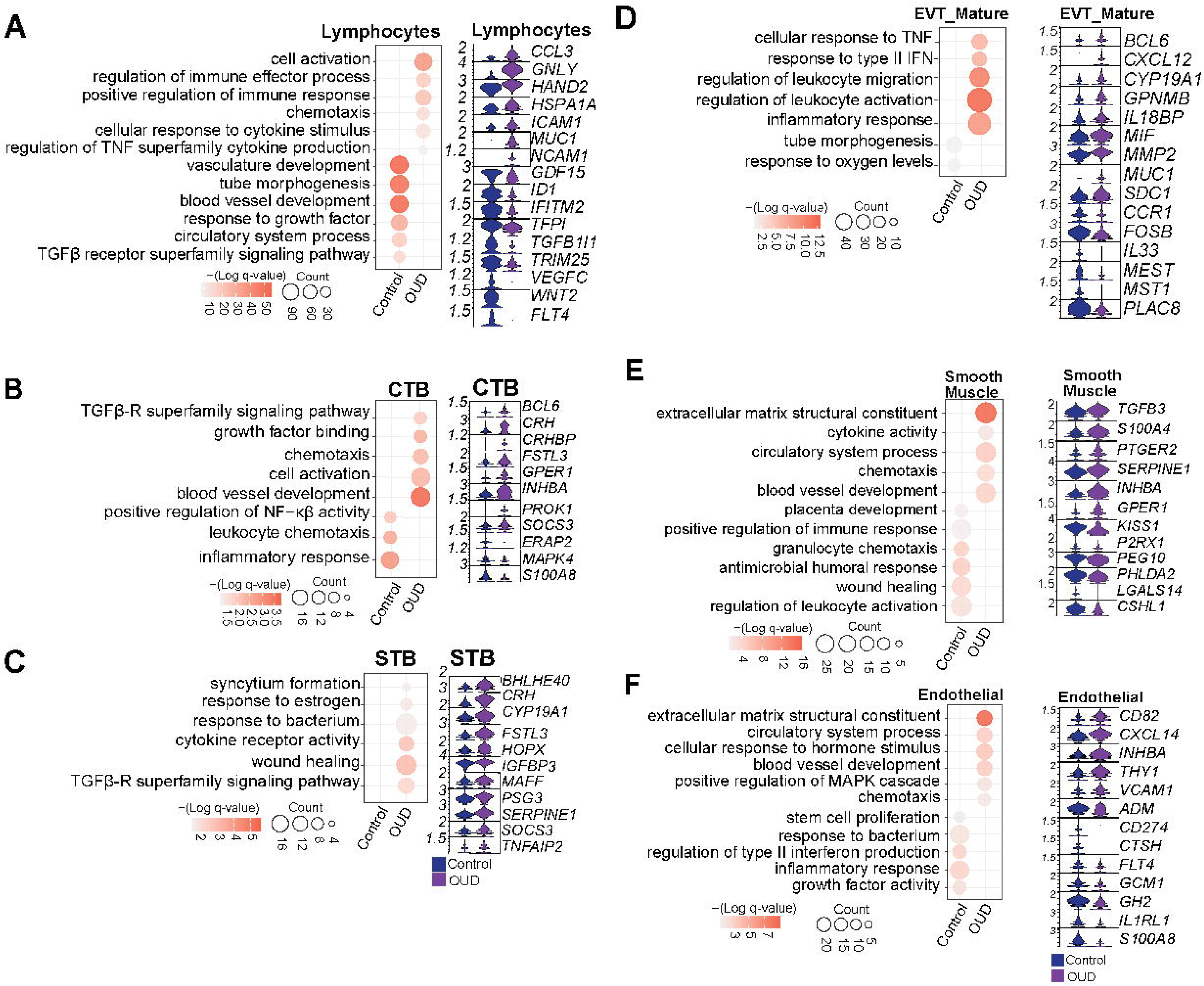
Maternal OUD±HCV impacts the placental transcriptional landscape at a spatial resolution. (A-F) Bubble plots of differentially expressed genes (DEGs) between control and OUD groups (left) and violin plots showing representative DEGs from the select GO terms (right) for the shown cluster. For the bubble plots, the size of the bubble represents the number of genes associated with GO term and the color represents the significance (q-value).

**Fig. S8:**
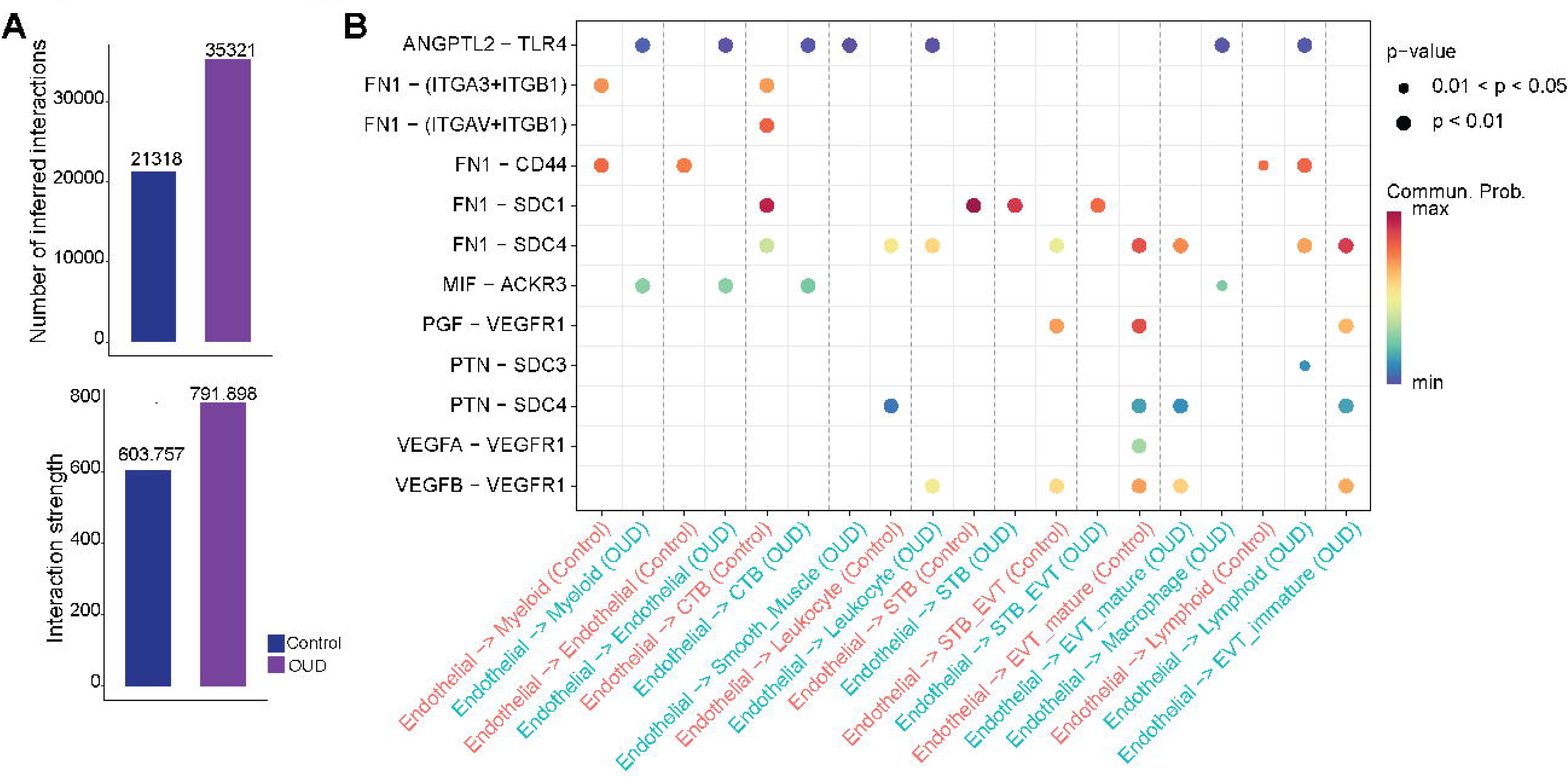
Maternal OUD impacts communication networks in the placenta. A) Bar plots depicting the overall number of ligand-receptor interactions (top) and interaction strengths (bottom) between groups. B) Bubble plot comparing the significant ligand-receptor pairs between control (left) and OUD (right) placenta, which contribute to the signaling from Endothelial cells to all other cell type clusters. Dot color reflects communication probabilities and dot size represents computed p-values. Empty space means the communication probability is zero.

## Notes

### Competing Interest Statement

The authors have declared no competing interest.

